# Transcriptional signatures associated with waterlogging stress responses and aerenchyma formation in barley root tissue

**DOI:** 10.1101/2023.10.06.561260

**Authors:** Orla L. Sherwood, Rory Burke, Jennifer O’Rourke, Conor V. Whelan, Frances Downey, Louise Ryan, Eoin F. McCabe, Zixia Huang, Carl K. Y. Ng, Paul F. McCabe, Joanna Kacprzyk

**Affiliations:** School of Biology and Environmental Science, University College Dublin, Dublin 4, Ireland

## Abstract

The negative impact of soil waterlogging on crop production is expected to increase due higher frequencies of extreme rainfall events arising from climate change. Consequently, understanding the molecular mechanisms that enable plants to mitigate waterlogging stress is critical for breeding programmes, particularly in the case of waterlogging-susceptible crop species, such as barley (*Hordeum vulgare*). Aerenchyma formation is a key morphological adaptation allowing plants to cope with waterlogging stress and hypoxic conditions, however, the genetic regulation of its development in barley remains largely unresolved. In this study, two barley cultivars with contrasting waterlogging tolerance (Franklin and Yerong) were subjected to waterlogging stress, followed by analysis of phenotypic traits including root aerenchyma formation, and transcriptomic profiling of root tissue samples. Differential expression analyses identified genes transcriptionally responsive to 24 and 72 h of waterlogging in both cultivars, and highlighted metabolic adaptations, regulation of ROS signalling and management of stress responses as key elements of the waterlogging response. The results revealed large intra-individual variations in root aerenchyma formation, and these variations were exploited to isolate 81 candidate aerenchyma-associated genes from the generated RNA-seq datasets. Network analyses suggest the involvement of the DNA damage response gene, *DRT100* and cell wall modifying genes, *XHT16* and *XHT15* as regulatory hub genes in aerenchyma formation. Collectively, the generated data provide insights into transcriptional signatures associated with the barley root responses to waterlogging and aerenchyma formation, informing our understanding of strategies that plants employ to cope with the negative impacts of heavy rainfall.

## Introduction

Waterlogging is a severe abiotic stress. Due to lower oxygen diffusion rates in water than in air, the oxygen availability to submerged plant tissues is reduced, leading to an average of 33% yield loss in affected crops (Tian et al., 2021). Altered rainfall patterns are consequences of climate change, resulting in an increased risk of flooding in many areas (Bailey-Serres et al., 2012; Westra et al., 2014). It has been projected that extreme precipitation events will occur 32% more frequently by 2100 (Thackeray et al., 2022); therefore, development of flood-tolerant crop cultivars is urgently required to protect food security. Plant tolerance to flooding stress differs widely across species: wetland plants such as rice generally exhibit high tolerance, while many other key crops, including barley (*Hordeum vulgare*) show sensitivity to even short-term waterlogging, further underscoring the need to develop tolerant varieties (de Castro et al., 2022; Smirnoff & Crawford, 1983). In barley, the focus species of this study, yield losses due to waterlogging can reach up to 70%, depending on the stress duration, soil type, temperature, and plant developmental stage when stress is imposed (Liu et al., 2020; de San Celedonio et al., 2016; Pang et al., 2004). As barley is the fourth most important cereal crop globally (Tricase et al., 2018), it is critical to elucidate the molecular mechanisms underlying waterlogging stress adaptation in this species, and in this way, inform development of tolerant germplasm. Indeed, there is extensive evidence for genetic regulation of waterlogging tolerance in plants, including barley, suggesting a high potential to breed for improved waterlogging resilience. For example, quantitative trait loci (QTL) analysis revealed four loci associated with barley waterlogging tolerance (Zhou, 2011), and subsequent QTL meta-analysis led to the identification of 58 putative waterlogging tolerance-associated genes (Zhang et al., 2017).

With most breeding efforts focused on shoot biomass and seed yield, it has been suggested that the roots of cereals are a potential key for a second green revolution (Gewin, 2010; Maqbool et al., 2022). This is particularly relevant in the context of waterlogging, where the root is the first organ to be affected, leading to a drastic reduction of root length and biomass, and decreased root to shoot ratio, impairing a crop’s ability to deal with adverse environmental conditions (Herzog et al., 2016; Malik et al., 2003). To counteract these effects, plants respond to waterlogging stress through altered metabolism and morphology. Under waterlogged conditions, plants must adapt to low oxygen availability, particularly affecting roots as subterranean organs (Sauter, 2013; Yamauchi et al., 2018). Consequently, plants must undergo extensive adaptations, involving switching from aerobic to fermentative metabolism, and modifications of organ structure to promote plant survival (Pan et al., 2021a).

Anatomical adaptations to waterlogging includes aerenchyma formation (Fagerstedt, 2010) and adventitious root development (Jackson & Armstrong, 1999). Aerenchyma, a key trait that promotes waterlogging tolerance (Mustroph, 2018), is a tissue composed of interconnected spaces, to enable movement of gases between shoots and roots, and consequently alleviates hypoxia in submerged plant tissues (Jackson & Armstrong, 1999). Indeed, faster aerenchyma formation in adventitious roots under waterlogging stress is associated with increased survival rates (Zhang et al., 2015) and marker trait associations for aerenchyma formation were previously detected in barley (Broughton et al., 2015; Manik et al., 2022; Zhang et al., 2016a). Inducible aerenchyma formed through programmed cell death is stimulated by ethylene during waterlogging, and is likely mediated by reactive oxygen species (ROS) (Sasidharan & Voesenek, 2015). Indeed, respiratory burst oxidases (RBOHs) promoted aerenchyma formation in response to waterlogging stress, and *RBOH* genes were induced by ethylene (Yamauchi et al., 2017; Yamauchi & Nakazono, 2022; Zhang et al., 2017). Adventitious roots formed under waterlogging stress conditions are rich in aerenchyma and often have an enhanced structural barrier to prevent radial oxygen loss, thereby promoting plant survival (Sauter, 2013; Visser & Voesenek, 2005). Despite these critical roles of root system adaptations to waterlogging stress survival, relatively little is known about the responses of root tissue to waterlogging at the molecular level in barley, and in particular about the regulation of aerenchyma formation. However, recent progress in this area has been observed in terms of characterization of gene expression changes and proteomic profiling associated with response to waterlogging in barley roots (Borrego-Benjumea et al., 2020; Luan et al., 2018a), facilitated by the availability of a chromosome-scale assembly of the barley genome (Mascher et al., 2017, 2021) and a high-resolution barley transcriptome (Coulter et al., 2022). For example, Borrego-Benjumea et al. (2020) carried out whole root tissue transcriptional profiling under waterlogging stress and a larger scale study examined waterlogging-induced root transcriptomic signatures in 21 barley cultivars that led to isolation of 98 core waterlogging response genes (Miricescu et al., 2023). Root transcriptome profiling (Luan et al., 2023), and combination of genome-wide association studies (GWAS) and root transcriptome profiling (Luan et al., 2022), also led to identification of several barley waterlogging associated genes that were further functionally validated in *Arabidopsis thaliana*.

In this study, we explored the transcriptional signatures associated with responses to waterlogging in barley root tissue, focusing specifically on aerenchyma formation. Two barley cultivars that were previously reported to exhibit contrasting waterlogging tolerance, Franklin (sensitive) and Yerong (tolerant) (Broughton et al., 2015; Li et al., 2008; Luan et al., 2018b; Xue et al., 2010; Zhang et al., 2015; Zhang et al., 2016a; Zhou, 2011), were subjected to waterlogging stress. Root aerenchyma formation, adventitious root growth, and shoot growth were monitored over the stress duration of 21 days. RNA sequencing was used to characterize gene expression changes in adventitious root tissue after 24 and 72 h of waterlogging treatment. Two roots were sampled per plant, and notably, we observed a high level of intra-individual variation in percentage of root aerenchyma at both control and waterlogging stress conditions in both cultivars. We exploited this phenotypic variation to identify genes putatively associated with aerenchyma formation in barley and to infer a gene regulatory network (GRN) underlying development of this trait. Collectively, our data not only further elucidated the transcriptional responses to waterlogging stress in barley root tissue, but also provided specific insights into the genetic regulators underlying aerenchyma formation, a key anatomical adaptation for waterlogging resilience.

## Materials and methods

### Plant material and growth conditions

Seeds of two barley (*Hordeum vulgare)* cultivars, Franklin and Yerong were obtained from Genebank of IPK Gatersleben. Franklin is a two-row winter malting barley accession (Code HOR 15356) while Yerong is a six-row spring barley accession (Code BCC 1709) (Li et al., 2008). Seeds were surface sterilised in 20% bleach for 10 min, rinsed at least 5 times in sterile distilled water (SDW) and placed on 85 mm Whatman grade 1 filter paper soaked with 5 ml SDW in 90 mm Petri dishes (six seeds per plate). Petri dishes were sealed with Leukopore tape and placed in the dark for 3 d at 4°C. Following cold stratification, seeds were placed at 20°C in the dark for 4 d for germination. Plastic pots (9×9×10 cm) were filled within 1 cm of the top of the pot with autoclaved (40 min, 121°C) vermiculite as the growth medium (Product Code VERM, National Agrochemical Distributors Limited, Dublin, Ireland). Vermiculite was chosen as a semi-structured hydroponics growth substrate to permit the rapid and straightforward collection of clean, undamaged root tissue for subsequent analyses. After 4 d of growth, one seedling was transferred per pot, 1 cm below the surface of vermiculite soaked with 0.25% Hoagland’s solution [pH 5.8] (Product Code H2395-10L, Sigma-Aldrich, Gillingham, UK). Trays were watered three times weekly from above and excess Hoagland’s solution removed after 30 min. Plant growth conditions were 20°C (constant temperature), 16 h light /8 h dark photoperiod (85 µmol m^-2^ s^-1^) under fluorescent lamps (Product Code Master TL5 HO 24W840, Philips, Amsterdam, The Netherlands).

### Waterlogging treatment

Plants were grown until the 3-leaf stage (Zadok’s stage 13) at 21 d under the growth conditions above. To initiate the waterlogging treatment, each pot was placed in 17×17 cm plastic bag to prevent drainage, and subsequently placed into an empty 9×9×10 cm plastic pot to hold the bag securely in place. The bags were then filled with 0.25% Hoagland’s solution [pH5.8] (Product Code H2395-10L, Sigma-Aldrich, Gillingham, UK) up to 1 cm above the growth substrate. This level of liquid medium was maintained for the duration of the waterlogging experiment. Control plants were maintained in the previous growth conditions detailed and were watered according to normal watering schedule. Plants were waterlogged as detailed in Figure 2A.

### Shoot and root measurements and root sampling

Shoot height (distance from the soil surface to the tip of the tallest leaf in the plant) was first measured directly prior to waterlogging treatment initiation, and at each sampled time point (day 1, 3, 7 and 21). For root phenotyping and sampling, plants were removed from the pots and the root system gently washed with water to remove vermiculite, followed by recording the number and length of all adventitious roots for day 1, 3 and 7. Due to high number of roots and difficulty in separating highly interwoven roots without causing damage these measurements were not taken after 21 days of treatment. Further, for quantification of aerenchyma (day 1, 3, 7 and 21) and transcriptomic profiling (day 1 and 3), two adventitious roots were sampled per plant. Sampled roots were at least 6 cm long. The 2 cm long root section from 1 to 3 cm away from the root tip was flash frozen in liquid nitrogen, and stored at −80°C until RNA extraction. Additionally, the 2 cm long root section from 3 to 5 cm away from the root tip was collected and stored in water for aerenchyma quantification. SPSS Version 27 (IBM Corp, 2020) was used for statistical analyses of phenotypic traits.

### Hand-sectioning of adventitious roots and aerenchyma quantification

Collected root segments were placed in individual vertically positioned moulds constructed from 2 ml syringes with tip cut-off, and embedded in 5% (w/v) liquid agar (Product Code A1296, Sigma-Aldrich, Gillingham, UK) (Figure S1). The embedded roots were manually sliced into thin sections (∼0.25 mm thick) using a double-edged razor blade (Solimo, Washingon, USA). The sections were kept in water, until imaging on microscope slides using a Leica DM500 light microscope connected to Leica MC170 HD camera (100 x magnification). For aerenchyma quantification images were analysed with ImageJ v1.50i (Schneider et al., 2012) to calculate overall cross section area of the root, number of aerenchyma pockets and aerenchyma area for each sample. SPSS Version 27 (IBM Corp, 2020) was used for statistical analyses of aerenchyma measurements.

### Root homogenisation, sample pooling, RNA extraction and sequencing

Frozen root tissue was manually homogenised in liquid nitrogen in 1.5 ml Eppendorf tubes using microtube pestles and 0.1 mm glass beads (Product Code 11079-101, Biospec Products, Inc., Oklahoma, USA). The pooling of individual samples of homogenised root sections prior to RNA extraction was informed by the previously performed aerenchyma quantification that revealed high intra-individual (root to root) variation in this trait. Briefly, for two adventitious roots collected from each plant; the root with lower aerenchyma percentage was categorised as L_AE, and the root with higher aerenchyma percentage as H_AE. For each time point (1 and 3 days), condition (control, waterlogging) and cultivar (Yerong, Franklin), the RNA was extracted separately from pooled H_AE and pooled L_AE homogenized root samples for each of 3 independent experimental repeats. Total RNA was extracted from the homogenised root tissue using the Qiagen RNeasy Plant Mini Kit (Product Code 74904, Maryland, USA) with an on-column DNase digestion step (RNase-Free DNase Set, Product Code 79254, Qiagen, Maryland, USA). Obtained RNA was quantified using an ND1000 Nanodrop (Thermo Scientific, Massachusetts, USA) and its integrity determined using an Agilent 2100 Bioanalyzer (Agilent). Library preparation and mRNA sequencing (NovaSeq PE150, 6G raw data per sample, poly-A selection) was carried out by Novogene (Cambridge, UK). In total, forty-eight RNAseq datasets were generated: three experimental repeats – two cultivars x two timepoints x two treatments x two aerenchyma categories (H_AE and L_AE) (Table S1).

### Differential gene expression analysis

The differential gene expression (DGE) analysis was performed using Galaxy’s graphical user interface for quality checking and trimming (https://usegalaxy.eu/; Afgan et al., 2018), through the command line for mapping and expression quantification and finally through R for differential gene expression analysis. First, the quality of raw reads was assessed with FastQC (v0.11.9) (Andrews, 2010) using default settings. The sliding-window function in Trimmomatic (v0.38.1) (Bolger et al., 2014) was used to trim windows of 4 bases below an average Phred score of less than 20 (99% accuracy of base call). Trimmed reads were mapped to the most comprehensive and resolved reference transcriptome available for barley, BaRTv2.18 (https://ics.hutton.ac.uk/barleyrtd/bart_v2_18.html; Coulter et al., 2022), through the command line with Salmon (v0.12.0) (Patro et al., 2017), using--validateMappings and –gcBias parameters. Raw expression counts of the transcripts based on BaTRv2 gene-transcript annotation were aggregated according to the gene-level counts using the R package *tximport* (Soneson et al., 2016). Subsequently, DGE analysis was carried out in RStudio (v2023.06.2) (RStudio Team, 2020) through R (R version 4.2.2) (R Core Team, 2022) using the package *DESeq2* (Love et al., 2014). Two different types of DGE analyses were performed. Firstly, to identify genes that might be associated with aerenchyma formation, the H_AE vs L_AE comparisons were performed for each time point, condition (waterlogging, control) and cultivar. Secondly, for characterization of the effect of waterlogging treatment itself on gene expression (WL vs CRT comparisons), the H_AE and L_AE datasets for each time point, condition and cultivar, were treated as technical replicates using DESeq2 function *collapseReplicates* (Table S1). In all analyses, the genes with less than 10 counts were removed, and Benjamini and Hochberg (Benjamini & Hochberg, 1995) adjusted p-value cut-off of 0.05 applied. Finally, the R package EnhancedVolcano (Blighe et al., 2018) with log _2_fold change cutoff set to 0.5 and *p*-value cutoff set to 10e^-3^ was used to generate volcano plots (Figure S2).

### Enrichment analyses

The BaRTv2 based annotations are not compatible with the available gene enrichment tools that are typically based on previous barley reference assemblies such as Morex V3 (Mascher et al., 2021). Therefore, the BaRTv2 annotations of genes isolated by DGE analyses were first converted to corresponding Morex V3 (Mascher et al., 2021) gene names using a file obtained from Dr. Linda Milne (Table S2) (Milne et al. 2021). This conversion of gene IDs currently provides corresponding MorexV3 IDs for ∼50% of BaRTv2 IDs. The gene enrichment analyses for identified (i) waterlogging responsive, and (ii) aerenchyma associated genes, were subsequently carried out on MorexV3 gene IDs using ShinyGO v0.75c (Ge et al., 2020). ShinyGO output was set to 30 GO terms with a false discovery rate (FDR) cut-off 0.05. Additionally, as the key focus of this study was identification of mechanisms underlying root aerenchyma formation and due loss of data in converted from BaRTv2 to MorexV3 IDs, another GO enrichment tool in PlantRegMap (2019-10-11) (Jin et al., 2015, 2017; Tian et al., 2020) was also used. *Arabidopsis thaliana* homologues were first identified using the PlantRegMap ID mapping tool by inputting BaRTv2 protein FASTA sequences of the 81 aerenchyma-associated genes, resulting in identification of 33 homologous *A. thaliana* genes. These 33 *Arabidopsis thaliana* homologues were used for the PlantRegMap GO term enrichment (*p-*value cut-off 0.01). Additionally, the motif-based Transcription Factor Enrichment function of PlantRegMap was used (*p*-value cut-off 0.01) to identify the putative upstream transcriptional regulators of this gene list.

### Gene regulatory network (GRN) construction

The GRN underlying aerenchyma formation was created using the GeneMANIA v3.5.2 (Warde-Farley et al., 2010) plug-in in Cytoscape v3.9.1 (Shannon et al., 2003). Arabidopsis *thaliana* homologues identified for isolated aerenchyma associated BaRTv2 barley genes were used as an input as the GeneMANIA tool is not compatible with the barley gene annotations. Addition of up to 20 predicted nodes was permitted. The GRN was clustered using clusterMaker2 v2.3.4 (Morris et al., 2011) with MCL (Markov Clustering Algorithm) clustering enabled and network granularity set to 4. Node size was set to correspond to degree of connectivity of nodes. Hub genes were identified based on nodes which were most connected in the network.

### Cross-study comparison

The comparison between results obtained here, and the universal barley 98 hypoxia responsive genes (MorexV3 annotation) recently reported by Miricescu et al. (2023) was performed. First, the available BaRTv2 gene IDs were retrieved for 98 hypoxia responsive genes from Miricescu et al. (2023) and GeneOverlaps package in R (Shen, 2023) was used to identify the genes that were also transcriptional responsive to waterlogging in this study.

### Data availability

RNA sequences and metadata for all root samples used in this publication has been uploaded to the NCBI Sequence Read Archive and are openly available under the Bioproject Accession number PRJNA956333 https://dataview.ncbi.nlm.nih.gov/object/PRJNA956333?reviewer=d98qm8ivmv96s40n46i69nu8cj.

## Results

### Root system adaptations mediating waterlogging tolerance in barley

Barley plants (cultivars Franklin and Yerong) were subjected to waterlogging treatment at 3 leaf stage and the shoot growth and root traits (aerenchyma formation, total length of adventitious roots and number of adventitious roots) were recorded at indicated time points (Figure 1). At day 1 and day 3 of waterlogging, segments of root tissue were also sampled for RNA isolation and sequencing (Figure 2A). The waterlogging treatment did not reduce the shoot growth rate over the 21 day treatment (Figure 1A), suggesting that plants have effectively activated responses conferring waterlogging tolerance over this period. Indeed, examination of percentage aerenchyma formation in the adventitious roots (two roots sampled per plant), suggested that waterlogging induced a significant increase in aerenchyma percentage in both cultivars, detected as early as after 1 day of treatment (Figure 1B). Generally, there was no difference in aerenchyma percentage between cultivars in this study’s experimental set up, apart from significantly higher (*p* = 0.017) percentage aerenchyma in adventitious roots of waterlogged Franklin (9.9%), compared to waterlogged Yerong (5.95%) after 1 day of treatment. This result may suggest faster initiation of aerenchyma formation in Franklin compared to Yerong. The total length of adventitious roots was not generally affected by waterlogging over a 7-day treatment period (Figure 1C), with the only significant difference observed for Franklin demonstrating higher total root length than Yerong under waterlogging conditions at day 7 (*p* = 0.016). The total number of adventitious roots, while not affected by waterlogging treatment itself over the investigated period, was lower for Yerong compared to Franklin at day 3 (*p* = 0.005) and day 7 (*p* <0.001) under waterlogging conditions (Figure 1D). Collectively, while both cultivars responded to waterlogging with immediate induction of aerenchyma formation in adventitious roots, the data also suggested a lower number of adventitious roots and lower total adventitious root length in Yerong under waterlogging conditions. This may indicate lower total volume of aerenchyma tissue available for gas exchange under waterlogging stress in this cultivar.

**Figure 1.**
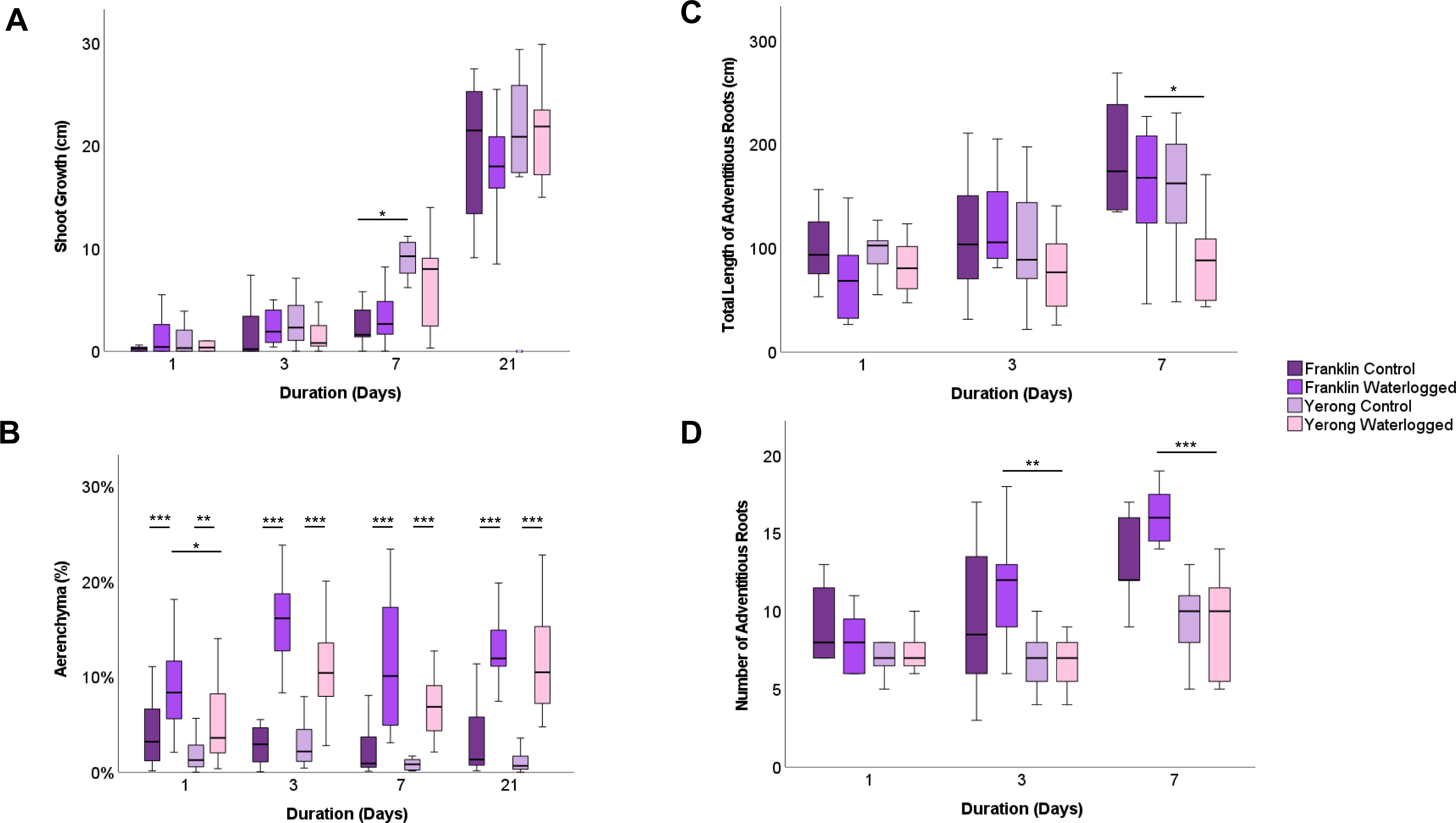
Response to waterlogging treatment observed in barley cultivars Franklin and Yerong. The waterlogging treatment was initiated at the 3-leaf stage. (**A)** Shoot growth was determined by measuring plant height prior to start of the treatment and after 1, 3, 7 and 21 d of waterlogging. **B**) Aerenchyma percentage in adventitious roots. Roots (2 per plant) were sampled 3-5 cm from the root tip and log10 transformed values were used for statistical analyses. **(C)** Mean total length of adventitious roots per plant. (**D**) Mean number of adventitious roots for each plant. Bars represent mean, error bars represent 95% CI. *** indicates *p* ≤ 0.001, ** indicates *p* ≤ 0.01, * indicates *p* ≤ 0.05 after 2-way ANOVA (*p*-value = 0.05) with Bonferroni correction for multiple testing using estimated marginal means in SPSS. Experiments were repeated three times with n = 12 cumulatively for all day 1 and 3 treatment groups, Day 7: Franklin Control n = 5, Franklin Waterlogged n = 8, Yerong Control n = 6, Yerong Waterlogged n = 7, Day 21: Franklin Control n = 9, Franklin Waterlogged n = 9, Yerong Control n = 10, Yerong Waterlogged n = 9 for **A**, **C** and **D**. For **B**, two roots were sampled from each individual plant, doubling n numbers for respective treatment durations i.e. n = 24 day 1 Franklin Control.

**Figure 2.**
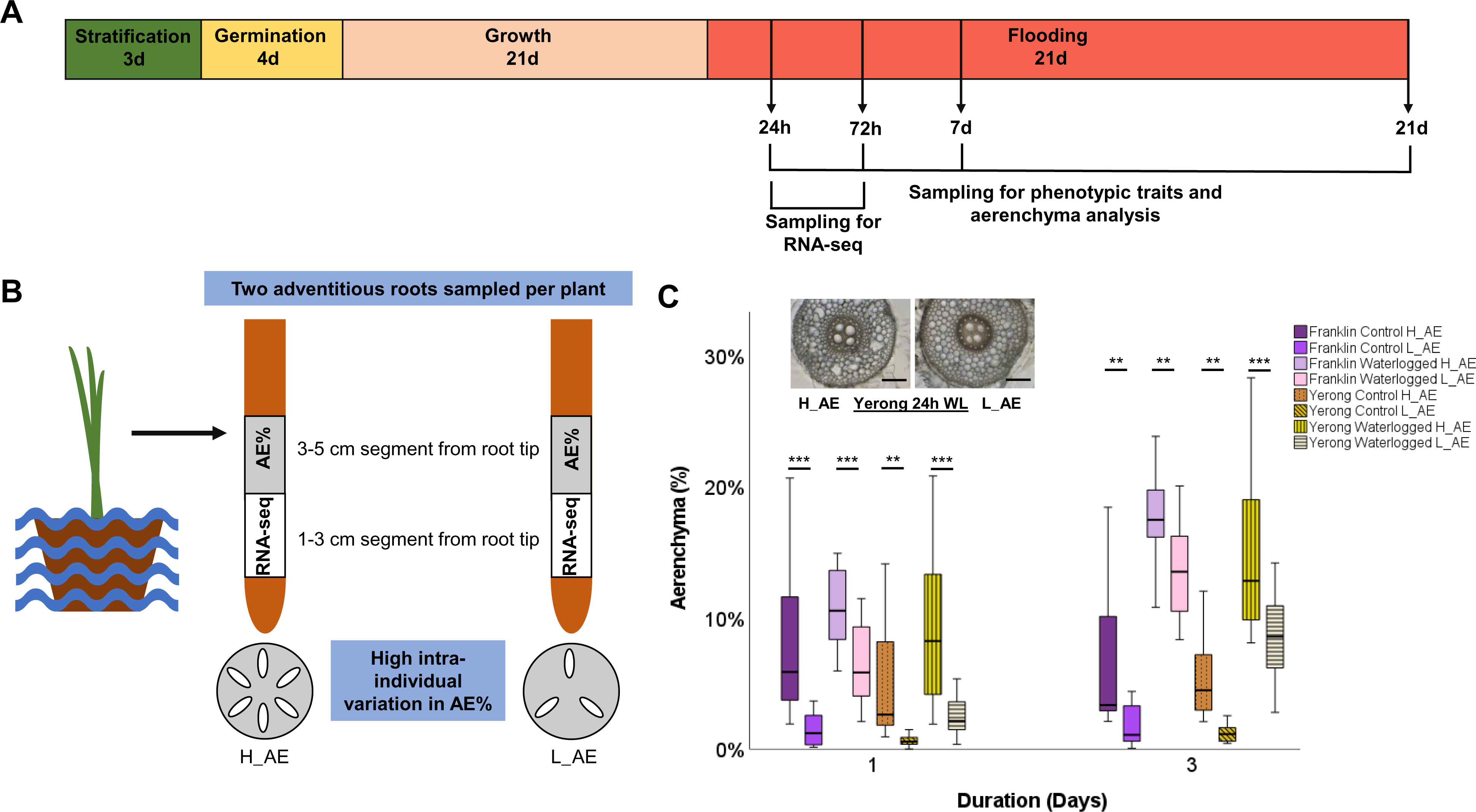
Timeline for RNA-seq sampling and observation of intra-individual variation in aerenchyma formation in adventitious roots. Barley plants were grown and sampled according to the timeline (**A**) where they were cold stratified for 3 d, transferred to 20°C in dark for germination for 4 d before planting in vermiculite soaked in 0.25% strength Hoagland’s solution. Plants were grown under 16 h light/ 8 h dark, 20°C constant temperature conditions for 21 d until plants reached the 3-leaf stage and were subsequently waterlogged with nutrient solution in plastic bags. Franklin and Yerong cultivars were sampled at 24 h and 72 h for RNA sequencing and for aerenchyma analysis and phenotypic traits at 1, 3, 7 and 21 d waterlogging treatment. Following 1 and 3 d treatment, root systems were washed in water and two roots were sampled from each plant for RNA sequencing (segment 1-3 cm from the root tip) and aerenchyma percentage analysis (segment 3-5 cm from the root tip) (**B**). For aerenchyma analysis, roots were embedded in liquid agar prior to hand-sectioning thin cross sections of root tissue and imaging using a light microscope at x100 magnification. Percentage aerenchyma was quantified for each root using ImageJ software. From each plant, both roots were compared and categorised as high (H_AE) or low (L_AE) aerenchyma forming roots relative to one another which led to the detection of high intra-individual aerenchyma percentage in roots sampled from the same plant (**B, C**). Graph indicates intra-individual variation in aerenchyma percentage of roots from the same plant, and provided images representing an example of intraindividual variation in % AE between two roots from the same waterlogged plant (cultivar Yerong, scale bar = 200 `µ`m). *** indicates *p* ≤ 0.001, ** indicates *p* ≤ 0.01 based on two-way repeated measures ANOVA (*p*-value = 0.05) with Bonferroni correction for multiple testing using estimated marginal means, error bars display 95% CI. Experiments were repeated three times n = 12 for each cultivar, treatment, timepoint and aerenchyma group.

### RNA sampling strategy to characterize waterlogging response in barley roots and isolate novel candidate aerenchyma associated genes

Root tissue was sampled for RNA sequencing after 1 and 3 days of waterlogging (Figure 2A,B) with 2 cm root segments (from 3 to 5 cm away from the root tip, directly above root segment used for determination of aerenchyma percentage) collected per sampled root. The aerenchyma percentage increased significantly in both tested cultivars after one day of treatment, and it was further enhanced after three days of waterlogging, suggesting that this sampling window is suitable for the capture of gene expression changes associated with aerenchyma formation. Furthermore, while in general there is a relatively limited number of datasets representing barley root response to waterlogging, the time points between 1 and 3 d were also used by several recently published studies exploring response to waterlogging (Borrego-Benjumea et al., 2020; Luan et al., 2023). Using the same timepoints therefore may facilitate comparisons between gene expression response to waterlogging observed across several datasets, different experimental set ups and cultivars. When analysing the root aerenchyma percentage, we also noted the high variability in aerenchyma percentage in the sampled adventitious roots observed for both cultivars (Figure 1B). Interestingly, the intra-individual variability in aerenchyma formation was substantial, thus offering an opportunity to isolate genes associated with aerenchyma formation by comparing gene expression profiles of root samples that differ in aerenchyma percentage within each condition and cultivar. We tested the extent of intra-individual variability in aerenchyma formation by measuring the aerenchyma percentage in the two adventitious roots collected from each plant; subsequently the root with lower aerenchyma percentage for each plant was labelled as L_AE and the root with higher aerenchyma percentage was labelled as H_AE (Figure 2B). The difference in aerenchyma percentage between L_AE and H_AE roots was significant at time points (day 1 and 3) for which tissue was sampled for RNA sequencing, under both control and waterlogging conditions, in both cultivars (Figure 2C). The intra-individual differences (Control: H_AE v L_AE, WL: H_AE v L_AE) and treatment induced differences (WL v C) in aerenchyma percentage for both cultivars and time points are presented in Table 1. To exploit the potential of this sizeable intra-individual variability in aerenchyma formation for identification of aerenchyma associated genes, during each independent experimental repeat, the RNA from pooled H_AE and pooled L_AE roots collected was sequenced separately for each time point (1 and 3 days), condition (control, waterlogging) and cultivar (Yerong, Franklin). Consequently, combining the H_AE and L_AE datasets facilitated determination of response to waterlogging stress for each timepoint and cultivar, whereas comparisons of matched H_AE and L_AE datasets enabled high resolution isolation of genes responsible for differences in aerenchyma formation.

**Table 1.** Increase in aerenchyma percentage resulting from waterlogging treatment (WL vs C) and intra-individual differences in aerenchyma percentage for each cultivar, treatment and timepoint. Barley plants were waterlogged for 24 h or 72 h and two roots were subsequently sampled from each plant for aerenchyma percentage quantification. For waterlogged (WL) vs control (C) comparisons, mean aerenchyma percentage was calculated for the two roots of the same plant prior to calculation of overall mean for each cultivar and timepoint. For high aerenchyma (H_AE) vs low aerenchyma (L_AE) comparisons, differences in aerenchyma percentage was calculated for both roots of the same plant prior to calculation of overall mean for each condition. Values represent mean ± SEM of three independent experimental repeats with n = 12 for each cultivar, treatment and timepoint. WL – waterlogged; C – control.

| Differences in % AE | 24 h |  |  | 72 h |  |  |
| --- | --- | --- | --- | --- | --- | --- |
| Cultivar | WL vs C | C: H_AE vs L_AE | WL:H_AE vs L_AE | WL vs C | C: H_AE vs L_AE | WL:H_AE vs L_AE |
| Franklin | 4.35± 0.55% | 6.76± 2.20% | 5.5± 1.65% | 10.1± 2.99% | 3.92± 1.10% | 4.21± 1.09% |
| Yerong | 2.74± 1.65% | 4.57± 2.00% | 6.75± 1.07% | 8.2± 0.54% | 4.59± 1.64% | 6.24± 1.85% |

### Transcriptional response to waterlogging stress in barley roots: pathways related to redox state management, metabolism and stress responses

Differential gene expression analysis was carried out to characterize the transcriptional response of barley roots to 24 h and 72 h of waterlogging stress (Figure 3A, Table S3A), using pooled H_AE and L_AE datasets for each timepoint, condition and cultivar. This revealed extensive changes in gene expression induced in both cultivars (Figure 3A, Table S3A). Further, we isolated cultivar specific- and core-DEGs, regulated in the same direction at both time points (‘general waterlogging response’, Figure 3B), or unique to only 24 h (early waterlogging response, Figure 3C) or 72 h (late waterlogging response, Figure 3D) time points. This revealed a greater overlap between the cultivars for general (32% overlap) and early (21% overlap) transcriptional response to waterlogging, compared to transcriptional changes unique to late (72 h) time points, where only ∼6% of DEGs were differentially regulated in both Yerong and Franklin. Subsequently, we investigated if differences in basal gene expression between the cultivars could explain the sizeable differences in waterlogging responses between Franklin and Yerong. However, while there were large differences in basal (control) gene expression between Franklin and Yerong at both 24 and 72 h, they provided putative explanation for only between 9 and 26 % of cultivar specific differences in gene expression in response to waterlogging (Figure S3). The transcriptional signature induced by waterlogging stress in this study was also compared to the list of core waterlogging-response genes recently identified by Miricescu et al. (2023), revealing a significant overlap and strong association between the response observed for both cultivars and at both time points (Table S4A), with the same direction of regulation (Table S4B). Collectively, generated data underscore that both core and cultivar specific responses to waterlogging stress exist and need to be further dissected to elucidate transcriptional signatures related to waterlogging tolerance and sensitivity.

**Figure 3.**
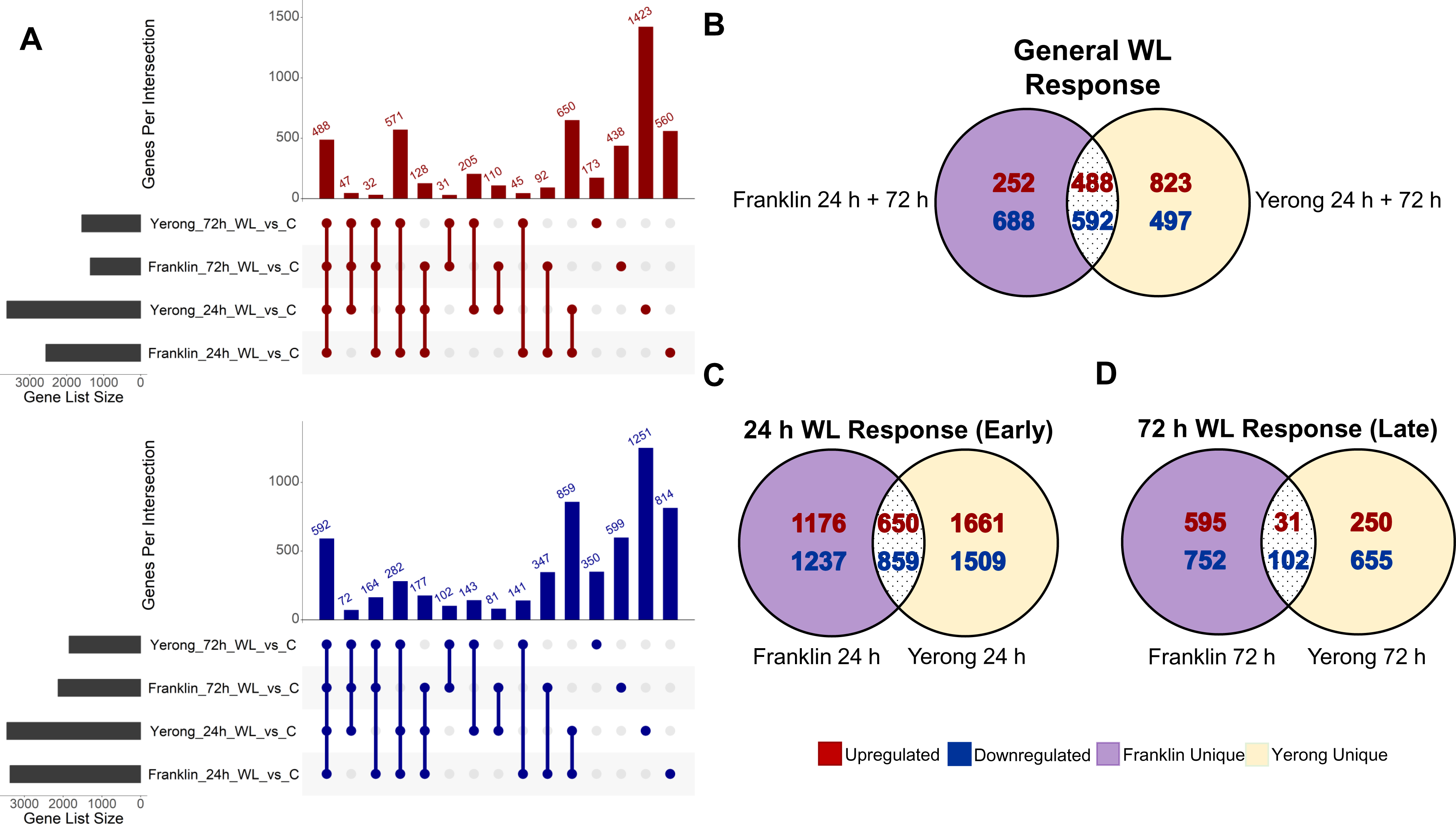
Transcriptional response to waterlogging stress in barley roots. **(A)** UpSet plots representing the overlapping DEGs upregulated (red) and downregulated (blue) in response to waterlogging (WL) at 24 h and 72 h treatment compared to respective control in each cultivar. Gene list size represents number of DEGs for each comparison and genes per intersection corresponds to the number of shared DEGs among the comparisons marked as connected dots below. The single dots represent DEGs unique to one comparison. Each comparison represented DEGs under waterlogging treatment with control as the baseline level. Venn diagram representing the general waterlogging responses. General waterlogging response genes are those commonly differentially regulated at point timepoints and in both cultivars (**B**). Number of DEGs unique to the 24 h (**C**) and 72 h (**D**) timepoint are shown in respective Venn diagrams indicating DEGs common to both cultivars and uniquely differentially expressed. Upregulated genes are shown in red; downregulated genes are shown in blue. Franklin unique expressed genes are highlighted in purple areas; Yerong unique genes are shown in yellow areas.

To further explore the core, and cultivar specific, responses to waterlogging stress and get insights into relevant biological pathways, gene ontology enrichment analysis was carried out on the isolated DEGs using ShinyGO (Ge et al., 2020) (Figure 4, S4, S5). As anticipated, the general (both time points) core (both cultivars) response to waterlogging resulted in enriched pathways involved in metabolism, stress mitigation and redox state, the pathways that are central components of stress responses in plants (Armstrong & Drew, 2002; Pan et al., 2021a; Teoh et al., 2022; Yamauchi et al., 2018). For example, the upregulated general core genes were enriched in glycolytic processes, carbohydrate catabolic process and oxidoreductase activity, as well as pathways associated with metabolism modulation and response to stress/stimuli, (e.g. “carbohydrate metabolic process”, “monocarboxylic acid metabolic process”, “response to chemical”) (Figure 4). The downregulated general core DEGs enriched oxidoreductase activity, catabolic processes, response to stimuli and transferase and transporter activity. This suggests that the general waterlogging response, independent of cultivar or stress duration, is comprised of mitigating stress through modulation of metabolism, activating stress responses and managing redox state to promote plant homeostasis.

**Figure 4.**
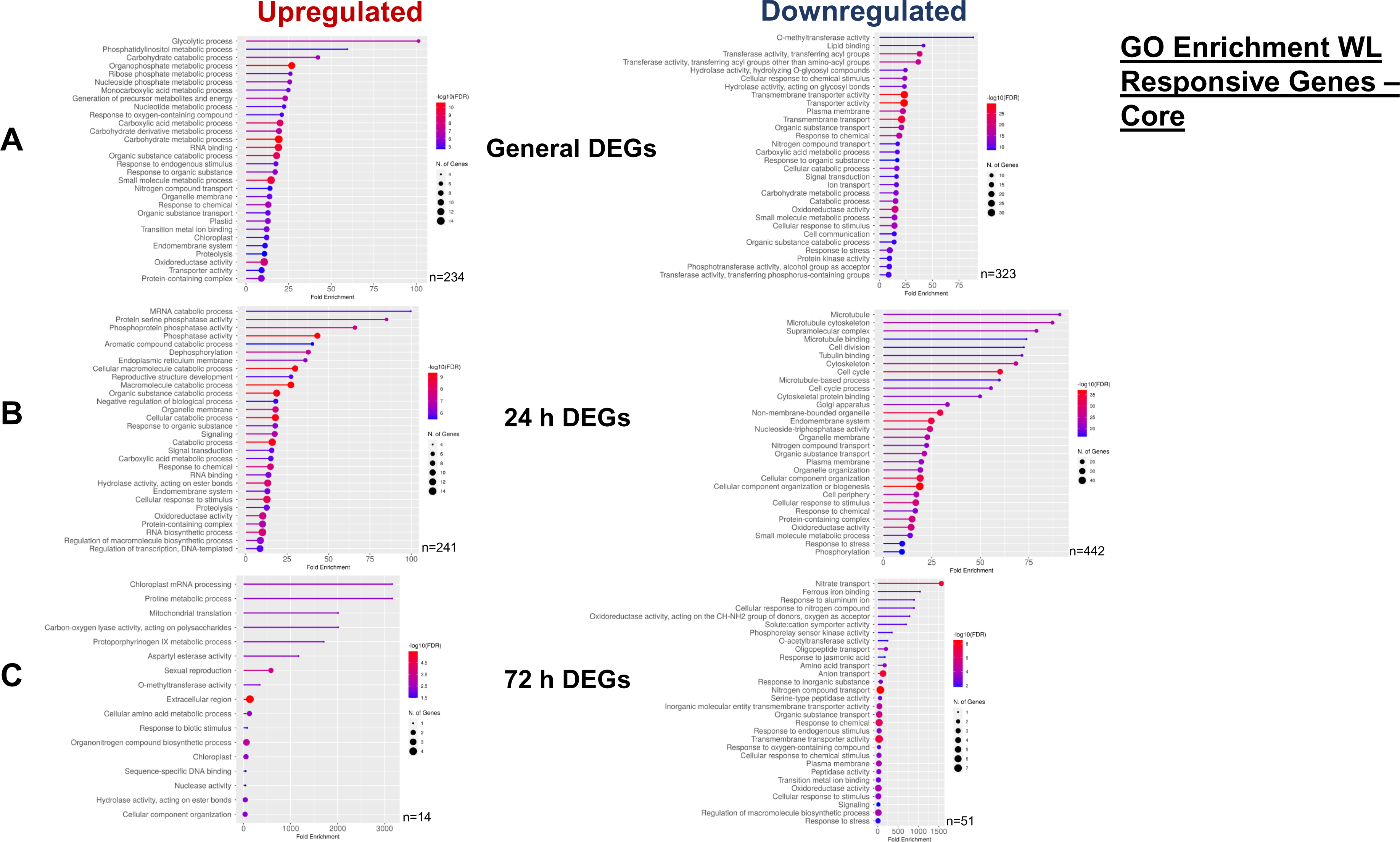
Gene ontology (GO) enrichment of WL response genes. Upregulated and downregulated DEGs associated with core (both cultivars), for general (both timepoints) (**A**), 24 h unique (**B**) and 72 h unique (**C**) timepoints were identified following DESeq2 analysis. BaRTv2 gene names were converted to MorexV3 gene names (represented by n for each GO analysis) and subsequently input into ShinyGO v0.75c. ShinyGO output was set to 30 GO terms and FDR cutoff as 0.05. The lollipop graph shows fold enrichment on the x-axis with size of dot representing number of genes associated with particular GO term. Colour of line and dot represents -log10(FDR).

The early (24 h) core waterlogging response up- and down-regulated DEGs reiterated the importance of redox regulation in waterlogging response (“oxidoreductase”) but also highlighted phosphorylation state-associated terms (e.g. “protein serine phosphatase activity”, “phosphoprotein phosphatase”, “dephosphorylation” and “phosphorylation”) and stimuli/stress response associated terms (e.g. “response to stress”, “response to chemical” and “cellular response to stimulus”), suggesting that modulation of phosphorylation state is associated with early stress response in barley. Interestingly, early (24 h) core waterlogging downregulated DEGs were enriched in terms including “cell division”, “cell cycle” and “cell cycle process” that may suggest early arrest of cell cycle during waterlogging stress. The later (72 h) specific core response upregulated DEGs highlighted GO terms related to hydrolase, transferase, lyase and esterase enzyme activities (e.g. “carbon-oxygen lyase activity, acting on polysaccharides”, “aspartyl esterase activity”, “O-methyltransferase activity” and “hydrolase activity, acting on ester bonds”) that may allude to enzymatic modulation of cell wall components during waterlogging. Furthermore, gene enrichment analysis of cultivar specific core, early (24 h) and (72 h) responses to waterlogging, suggested that at pathway level the gene expression changes observed in Yerong and Franklin are generally more similar than at gene list level (Figure S4, S5) highlighting terms associated with oxidoreductase activity, catabolic processes, cell wall modification and responses to stress/stimuli. However, the GO term enrichment suggested that there might be some differences in terms of timing of cultivar specific responses. For example, the terms related to oxidative stress (e.g. “reactive oxygen species metabolic process” and “response to oxidative stress”), phosphorylation state (e.g. “protein phosphorylation”, “phosphotransferase activity, alcohol group as acceptor”, “protein kinase activity” and “pyrophosphatase activity”) and cell wall adaptation (e.g. “transferase activity, transferring phosphorus-containing groups”, “transferase activity, transferring glycosyl groups” and “peptidase activity”) were enriched by early (24 h) unique DEGs specific to Yerong but not to Franklin cultivar. While for 72 h unique genes, the enrichment of these terms is observed for both cultivars, suggesting more transcriptional reprogramming in Yerong for these terms at the earlier timepoint.

### Isolation of aerenchyma associated genes and constructing gene regulatory networks underlying aerenchyma formation in barley

The observed high intra-individual variability in aerenchyma percentage (Table 1, Figure 2C) offered a unique opportunity to identify genes putatively associated with aerenchyma formation in barley roots due to separately sequencing the RNA sampled from high aerenchyma (H_AE) and low aerenchyma (L_AE) roots derived from the same individuals, that for each experimental repeat were pooled for each time point (24 h and 72 h), condition (control, waterlogging) and cultivar (Yerong, Franklin). Differential gene expression analysis was carried out to compare the transcriptional profile of H_AE and L_AE samples at each time point, condition and cultivar (8 comparisons in total, Figure 5A). The 81 DEGs (annotation based on BaRTv2 transcriptome) that were present in at least two of H_AE vs L_AE comparisons were considered to be putatively associated with aerenchyma formation and used for further analyses (Table S3B). Among these aerenchyma responsive genes were three peroxidases, two kinases (one calcium dependent), two glycosyltransferases and a proteolytic enzyme bromelain. Moreover, 19 of the 81 isolated DEGs were differentially regulated under waterlogging conditions at least one time point, for at least one cultivar (Table S3C), and 14 of these 19 genes were differentially regulated at the 24 h waterlogging treatment timepoint for at least one cultivar. Some of the most interesting candidates from this aerenchyma associated, waterlogging induced gene list include peroxidases, which have been implicated in positive regulation of programmed cell death responses in plants (Graças et al., 2020, Zhang et al., 2022) and an S-type anion channel SLAH3 (encoded by *BaRT2v18chr1HG047880*), that was previously associated with plant immune responses (Liu et al., 2019).

**Figure 5.**
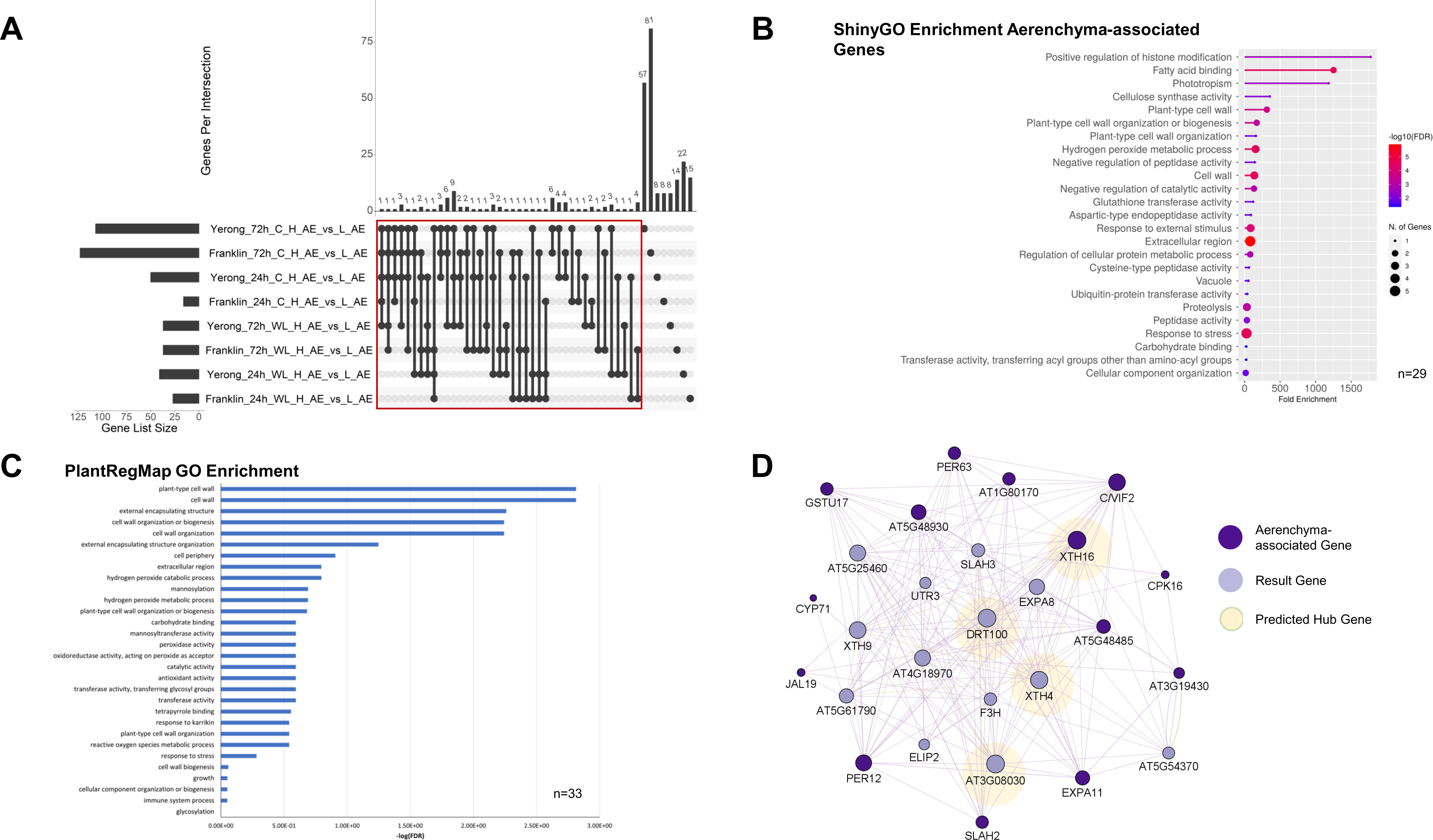
Aerenchyma-associated transcriptomic response in Franklin and Yerong barley cultivars to waterlogging stress. Differential gene expression analysis was carried out for high percentage aerenchyma (H_AE) vs low percentage aerenchyma (L_AE) comparisons in both treatments (waterlogging or control), both cultivars (Franklin or Yerong) and both timepoints (24 h or 72 h) with results summarised in UpSet plot (**A**). Gene list size represents number of DEGs for each sample for each comparison and genes per intersection represents number of overlapping DEGs between specific comparisons as indicated by the connected dots below. Single dots represent unique DEGs to one comparison. Each comparison represents DEGs in high percentage aerenchyma forming sample with low percentage aerenchyma sample as the baseline level. Red box indicates aerenchyma-associated DEGs common to at least two out of eight of the high vs low percentage aerenchyma comparison groups (81 DEGs). GO enrichment of aerenchyma associated DEGs (**B, C**). Aerenchyma-associated DEGs (BaRTv2 gene format) were converted to MorexV3 gene name where possible with n indicating number of converted genes to MorexV3 format. MorexV3 gene names subsequently input ShinyGO v0.75c for GO enrichment. Output was set to 30 GO terms and FDR cutoff as 0.05 (**B**). Aerenchyma-associated DEGs were input as BaRTv2 protein FASTA sequences into the ID mapping tool (BLASTp) in PlantRegMap to obtain *Arabidopsis thaliana* IDs, n represents number of DEGs which had a corresponding *A. thaliana* ID. *A. thaliana* IDs were input into the GO term enrichment tool in PlantRegMap (**C**) with p-value cut off of 0.01. Clustered gene regulatory network (GRN) of aerenchyma-associated genes with predicted hub genes (**D**). Aerenchyma associated genes (dark purple) with corresponding Arabidopsis gene IDs (n=33) were input into Cytoscape using the GeneMANIA v3.5.2 plugin where 20 result (light purple) nodes were inserted based predicted gene interactions. Node size corresponds to degree of connectivity of nodes with larger nodes more connected. Hub genes are identified by nodes with greatest connectivity and circled in green. The GeneMANIA network was clustered using the Cytoscape plugin clusterMaker2 v2.3.4with MCL clustering, granularity parameter set to 4. Interaction links in purple are predicted based on gene co-expression, in blue are co-localised genes and yellow are genes with shared protein domains. The edge thickness in the GRN corresponds to normalised max weight.

Further, to get pathway level insights into regulation of aerenchyma formation in barley roots, enrichment analysis was performed using ShinyGO (Figure 5B) and PlantRegMap (Figure 5C) tools. For enrichment analysis performed using ShinyGO, the BaRTv2 annotation had to be converted for MorexV3 gene names, that were available for 29 of 81 aerenchyma associated DEGs in BaRTv2 format. For enrichment analysis performed using PlantRegMap, the ID mapping function was used to identify corresponding *A. thaliana* gene names using BaRTv2 protein FASTA sequences of 81 aerenchyma-associated genes, resulting in identification of 33 homologous *A. thaliana* genes that were used for the PlantRegMap GO term enrichment. These GO term enrichment analyses (Figure 5B,5C) highlighted pathways associated with cell wall organisation/modification (e.g. “plant-type cell wall organisation or biogenesis”, “plant-type cell wall”, “cell wall organisation”, “cell wall”, “cell wall biogenesis” and “cellulose synthase activity”), stress responses (e.g. “response to external stimulus”, “response to stress” and “response to karrikin”), and redox state management (“glutathione transferase activity”, “hydrogen peroxide metabolic process”, “peroxidase activity”, “antioxidant activity”, “oxidoreductase activity, acting on peroxide as acceptor”, “hydrogen peroxide metabolic process”, “hydrogen peroxide catabolic process” and “reactive oxygen species metabolic process”), and regulation of plant immunity (“immune system process”). This provided validation of the isolated gene list of aerenchyma associated genes, considering that lysigenous aerenchyma formation involves programmed cell death of root cortical cells, and is a process associated with cell wall modifications (Gunawardena et al., 2001a; Saab & Sachs, 1996), management of ROS signalling (Ni et al., 2019; Pan et al., 2021b; Pan et al., 2022; Yamauchi et al., 2017) and modulation of immune signalling pathways (Karpinski et al., 2013; Mühlenbock et al., 2007). Interestingly, several enriched terms suggested also less explored pathways that may play a role in aerenchyma formation, or PCD in general, and should be further investigated in this context, including regulation of histone modification and response to karrikins as stress responsive compounds.

Subsequently, to further elucidate the genetic regulation of aerenchyma formation and isolate genes likely to play a central role in this process, we constructed a gene regulatory network (GRN) using Cytoscape (Shannon et al., 2003) and the GeneMANIA (Warde-Farley et al., 2010) plugin with 33 identified *A. thaliana* homologues of aerenchyma associated genes as an input (Figure 5C). GeneMANIA inserted 20 additional interactors, and the constructed GRN was clustered using the clusterMaker2 app (Morris et al., 2011), leading to isolation of a cluster of highly connected nodes (Figure 5D). The four most connected (≥20 connections) genes from this cluster were classified as hub genes, as they can play a central regulatory role in the constructed network and therefore warrant further investigation of their role in the context of aerenchyma formation. These hub genes were *DRT100* (DNA-DAMAGE REPAIR/TOLERATION 100) which plays a role in DNA damage repair and tolerance (Ghosh et al., 2013), *XTH16* (XYLOGLUCAN ENDOTRANSGLUCOSYLASE/HYDROLASE 16), *AT3G08030* (encoding a cell wall protein DUF642) and *XTH4* (XYLOGLUCAN ENDOTRANSGLUCOSYLASE/HYDROLASE 4). Finally, we used the PlantRegMap tool to identify putative upstream transcriptional regulators of the aerenchyma associated genes. The 33 identified *A. thaliana* homologues of aerenchyma associated genes were used as an input, and this analysis returned 39 transcription factors with significantly overrepresented targets (Table S5). Notably, many of the identified transcription factors were previously linked to stress responses (including water stress), plant immunity and PCD; with the majority of them (30) belonging to the ethylene response factor (ERF) family. Ethylene is a critical signal transducer that plays a central role in plant adaptation responses to low oxygen, including induction of aerenchyma (Gunawardena et al., 2001b; He et al., 1996; Ni et al., 2019; Yukiyoshi & Karahara, 2014) in addition to adventitious root formation (Negi et al., 2010; Qi et al., 2019; Visser et al., 1996), and indeed numerous ERFs were previously reported to mediate these responses (Hattori et al., 2009; Kitomi et al., 2011; Licausi et al., 2010; Trupiano et al., 2013; Yang, 2014).

## Discussion

Barley is a waterlogging-susceptible cereal crop. Waterlogging induced yield losses in this species vary from 10 to 50%, depending on the stress duration, plant developmental stage, soil type and temperature (Pang et al., 2004; Setter et al., 1999; Setter & Waters, 2003). Increased understanding of the genetic regulation underlying waterlogging tolerance mechanisms in barley is therefore urgently required to inform breeding programmes and to support development of climate-proofed, more waterlogging-tolerant, barley cultivars. The root system, as the first line of defence against soil waterlogging, undergoes metabolic and anatomical adaptations supporting survival of the whole plant, and therefore should be studied as a priority when early waterlogging stress responses are considered (Herzog et al., 2016; Langan et al., 2022). However, relatively limited resources are currently available in terms of transcriptomic (Borrego-Benjumea et al., 2020; Luan et al., 2023), proteomic or metabolomic datasets (Andrzejczak et al., 2020; Luan et al., 2018a; Luan et al., 2022) characterizing responses of barley root tissue to waterlogging stress or hypoxia. Here, we used a combination of plant phenotyping and transcriptomics (RNA-seq) to further our understanding of waterlogging responses in barely root tissue, with particular focus on elucidation of pathways and genes modulating formation of root aerenchyma, a key anatomical adaptation for waterlogging tolerance (Mustroph, 2018). The two barley cultivars used in this study were selected due to previous reports on their contrasting tolerance to waterlogging, with Yerong generally being reported to be more waterlogging tolerant and demonstrating more extensive aerenchyma formation compared to Franklin (Broughton et al., 2015; Li et al., 2008; Luan et al., 2018b; Xue et al., 2010; Zhang et al., 2015; Zhang et al., 2016a; Zhou, 2011). However, in this study, both cultivars appeared to cope well with applied waterlogging stress with no shoot growth reduction observed after 21 days of waterlogging. This can be explained by observed fast induction of aerenchyma in root tissue, allowing plants to thrive under hypoxic conditions. The percentage of root aerenchyma increased following 24 h of waterlogging in both cultivars, in line with other studies reporting development of this trait over a similar stress period (Manik et al., 2022; Teoh et al., 2022; Zhang et al., 2015; Zhang et al., 2016a). However, apart from a small, but significant difference of 3.95% in percentage aerenchyma between Franklin and Yerong after 24 h of waterlogging, that could be potentially due to slightly faster formation of aerenchyma in Franklin roots, the aerenchyma levels appeared similar in both cultivars over the investigated period. Furthermore, despite previous reports on waterlogging promoting formation of adventitious roots (Chen et al., 2002; Koramutla et al., 2022; Qi et al., 2019; Visser et al., 1996), we did not observe differences in total length and number of ARs over the investigated time period as a result of waterlogging treatment applied within each cultivar. However, under waterlogging conditions, the number of ARs (at day 3 and 7) and total root length (at day 7) were lower in Yerong compared to Franklin. These observations contrast with previous results for these cultivars, with, for example, Luan et al. (2018b) reporting an increase in both total AR length and number in waterlogged compared to control treatments in Yerong, but not in Franklin. The general lack of superior performance of Yerong, compared to Franklin under waterlogging conditions in this study, underscores that ranking barley cultivar as tolerant or sensitive depends on the exact experimental set up used (type of growth substrate, time/duration of treatment application, temperature, *etc*.) which makes it difficult to cross-compare the results (Miricescu et al., 2021). Indeed, neither cultivar tested here showed a reduction in shoot growth under waterlogging conditions over the investigated period, suggesting successful adaptation to the applied stress (likely due to observed fast development of aerenchyma in root tissue); thus, the experimental set up utilised, offered us an opportunity to characterize transcriptional responses in root tissue associated with effective mitigation of waterlogging stress on plant growth.

### Core and cultivar specific transcriptional response to wateorlgging in barley roots

Analysis of RNA-seq datasets, generated from collected root tissue, facilitated identification of core- and cultivar specific-genes differentially expressed in response to 24 h and 72 h of waterlogging. The transcriptomic analyses were performed using a highly resolved barley reference transcriptome that allows accurate transcript-specific RNA-seq quantification (BaRTv2) (Coulter et al., 2022). It needs to be highlighted that downstream analyses were challenging due to differences in gene identifiers between transcriptomic (BaRTv2) and genomic (MorexV3) resources for barley; for example gene enrichment analysis tool ShinyGO, used in this study, is compatible only with the MorexV3 gene IDs, requiring conversion of BaRTv2 identifiers to MorexV3 that in our case reduced the number of DEGs by ∼50% and likely resulted in loss of information depth. The gene expression changes observed at both time points, in both cultivars, highlighted GO terms associated with redox state management, metabolic adaptations and stress responses, suggesting that these processes play a key role in waterlogging response in barley (Figure 6). These GO terms have previously been linked to waterlogging responses in barley in other transcriptomic analyses (Borrego-Benjumea et al., 2020; Luan et al., 2023; Miricescu et al., 2023). In the first global transcriptomic study of barley root tissue under waterlogging stress, Borrego-Benjumea et al. (2020) similarly identified enriched GO terms in the upregulated gene set for metabolism (carbohydrate, glycolytic and nucleoside), general enrichment for “oxidoreductase activity” and “response to chemical” related GO terms in Yerong root tissue subjected to 72 h waterlogging stress. Likewise, short term waterlogging stress in Franklin and TX9425 barley cultivars, revealed higher representation of terms in metabolism modulation, catalytic activity and oxidation-reduction processes (Luan et al., 2023). This, together with observed significant overlap with core transcriptional response to waterlogging defined for barley by Miricescu et al. (2023) (Table S4A, 4B), places this research within the established literature on general transcriptomic responses in barley to waterlogging.

**Figure 6.**
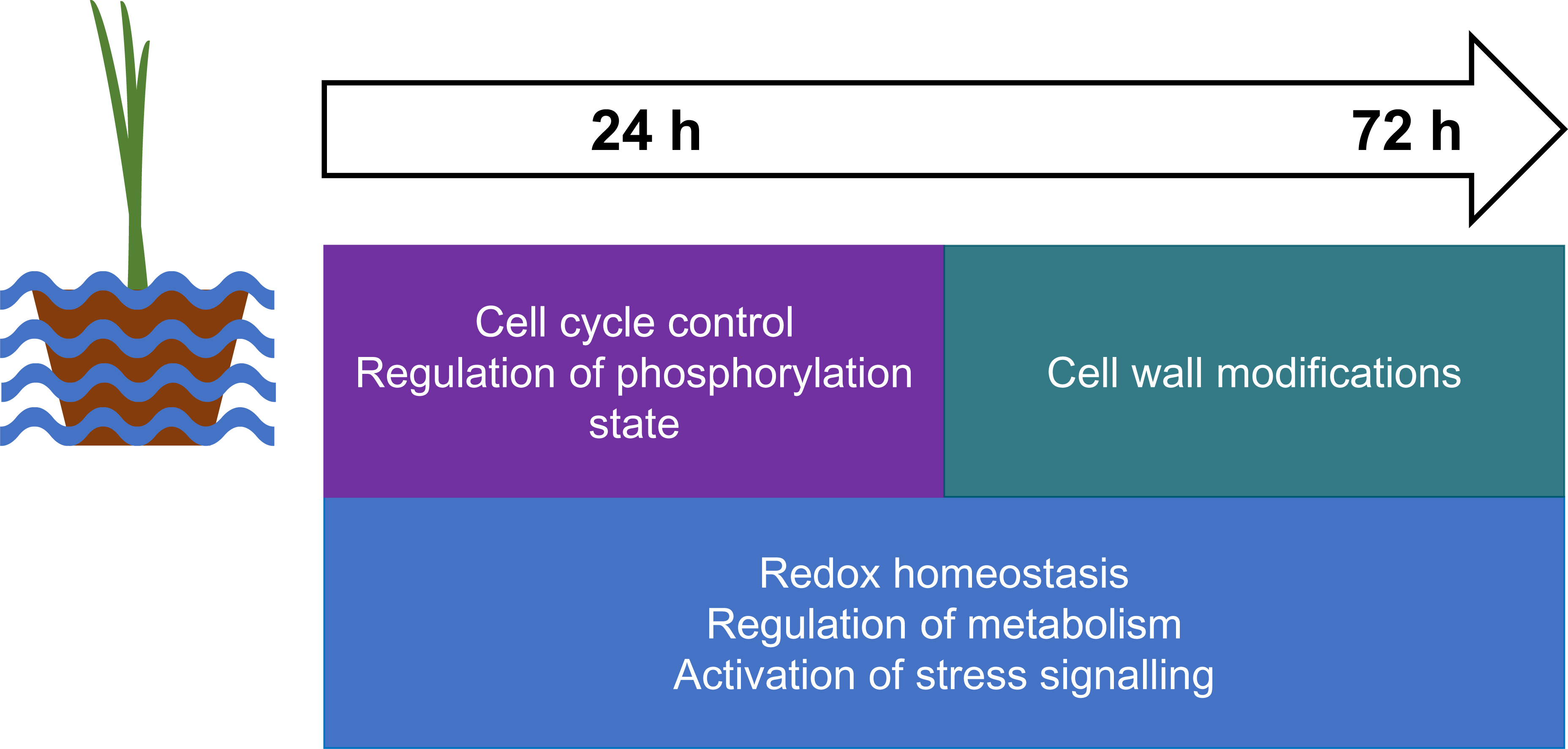
Summary of transcriptional changes in barley roots under waterlogging stress.

Gene enrichment analysis of genes differentially expressed in both cultivars and specific to the 24 h time point further underscored the importance of redox state management, but also highlighted that pathways playing a role in control of phosphorylation state and cell division contribute to early response to waterlogging stress. While their exact role in waterlogging responses is yet unresolved, kinases and modulation of phosphorylation state plays a role in many cellular reactions, including regulation of the activity of the hypoxia responsive protein respiratory burst oxidase H (RBOHH) in rice roots, that was linked to aerenchyma formation (Yamauchi et al., 2017). Kinase activity was also linked to managing glycolytic activity under anoxia via regulation of hexokinases in maize root tips (Bouny & Saglio, 1996). Further, the 24 h unique downregulated genes here were enriched in cell cycle related GO terms may underscore the requirement to manage cell division and cycle processes under stress. In fact, disruption of the cell cycle is associated with stress in Arabidopsis (Takahashi et al., 2019) and programmed cell death responses (Burke et al., 2023; Chen et al., 2017; Takahashi et al., 2019, Zebell & Dong, 2015). As plants actively respond to stress, cell cycle arrest is plausible, which minimises use of energy and resources and is potentially related to localized induction of programmed cell death and subsequent aerenchyma formation. In contrast, gene expression changes unique to the latter, 72 h time point, highlighted modifications of cell wall components occurring at this stage (Figure 6). This is supported by observations from other studies, that suggest that morphological changes associated with cell wall structure generally appear after an extended treatment duration (Watanabe et al., 2013), including the cell wall modifications after prolonged hypoxia treatment in maize (Gunawardena et al., 2001a), and suberin deposition observed only after long term waterlogging stress in rice (Watanabe et al., 2013) and under prolonged nutrient deficiency in barley (Chen et al., 2019).

### Genetic Regulation of Aerenchyma Formation in Barley

Previous studies reported that aerenchyma percentage varies substantially depending on the sampling location within the root (distance from the base or root) (Bouranis et al., 2006; Chen et al., 2021; Xu et al., 2013) but they did not investigate differences in aerenchyma levels observed between roots of the same individual. In this study, we observed a high intra-individual variation in this trait, demonstrated by large differences in percentage aerenchyma between two adventitious roots sampled per plant. Our data underscore that multiple roots should be sampled per plant to achieve an accurate representation of aerenchyma levels observed for particular genotype, under specific growth and treatment conditions. We exploited the observed high intra-individual variation in levels of aerenchyma for identification of putative aerenchyma associated genes through comparisons of transcriptional profiles of roots differing in percentage aerenchyma, sampled from the same individuals within each genotype, timepoint and treatment.

Using this approach, we isolated 81 candidate aerenchyma-associated genes (based on BaRTv2 transcriptome annotation). This list included genes encoding proteins previously reported in context of aerenchyma formation in range of different plant species, such as three peroxidases, two kinases (calcium-dependent protein kinase 18 and a putative serine/threonine-protein kinase-like protein CCR3), numerous carbohydrate modification associated proteins including a pectinesterase inhibitor 28 protein, xyloglucan endotransglucosylase/hydrolase, expansin, glycosyltransferase family protein 2 and polygalacturonase. This is in line with previous findings of peroxidases being upregulated under hypoxia stress (Hofmann et al., 2020; Mühlenbock et al., 2007) and during PCD regulation (Graças et al., 2020, Zhang et al., 2022), cell wall modifications being an important stage of the aerenchyma formation process (Gunawardena et al., 2001a; Leite et al., 2017; Pegg et al., 2020) and reported involvement of kinases in aerenchyma regulation (Wany et al., 2017; Yamauchi et al., 2017). A particularly interesting aerenchyma-associated candidate gene, that was also strongly induced by 24 h waterlogging in both cultivars (log _2_ fold change 20.81105832 in Franklin, 38.12043617 in Yerong), encodes an S-type anion channel SLAH3 (SLOW ANION CHANEL1 HOMOLOG3). In Arabidopsis SLAH3 was linked to immune responses (Liu et al., 2019), control of rapid stomatal closure (Zhang et al., 2016b), plasma membrane potential (Liu et al., 2023) and regulation of apoplastic reactive oxygen species-induced cell death (Jalakas et al., 2021). Further, SLAH3 promotes plant submergence stress responses in Arabidopsis by sensing hypoxia induced cytosolic acidification and mediating root anion efflux (Lehmann et al., 2021). The role of SLAH3 in regulation of cell death mediating aerenchyma formation, potentiated by hypoxic conditions, is therefore plausible.

The gene enrichment analysis further validated the list of aerenchyma-associated candidate genes, highlighting terms linked to processes previously reported to be involved in development of this trait. Gene enrichment analyses performed using (i) MorexV3 genome IDs and ShinyGo, and (ii) best Arabidopsis hits of identified aerenchyma associated genes and PlantRegMap tool, both pointed to cell wall modulation, redox state management and regulation of stress and immune responses, all of which were previously studied in context of aerenchyma formation in range of plant species (Armstrong et al., 2019; Basu et al., 2020; Chen et al., 2016; Gunawardena et al., 2001a; He et al., 2019; Leite et al., 2017; Mühlenbock et al., 2007; Ni et al., 2019; Pan et al., 2022; Pegg et al., 2020; Pucciariello & Perata, 2017; Rajhi et al., 2011; Saab & Sachs, 1996; Steffens et al., 2011; Wany et al., 2017; Wany & Gupta, 2018; Xu et al., 2013; Yamauchi et al., 2014, 2017). The functional enrichment analysis also highlighted that histone modifications may play a role in aerenchyma formation. This is in line with a study on wheat seminal roots, where blocking histone acetylation was found to impact waterlogging tolerance and aerenchyma formation through changes to cell wall degradation (Li et al., 2019). Finally, the “response to karrikins”, small organic compounds that are released during wildfires from burning plant material (Flematti et al., 2015), was among terms enriched among isolated aerenchyma associated genes. A recent study examined the impact of smoke solution on wheat seedlings under waterlogging stress (Komatsu et al., 2022). The karrikin-containing smoke solution alleviated the effect of flood stress on leaf growth, altered metabolic signatures associated with amino acids and maintained abundances of photosynthesis-related proteins in wheat plants under flooding stress. The role of karrikin signalling pathway is also emerging in relation to plant responses to other abiotic stress conditions, such as heat, cold, salinity and osmotic stress (Abdelrahman et al., 2022, Wang et al., 2018) and karrikin signalling was found to promote ethylene synthesis to modulate root architecture (Swarbreck, 2021). Considering the known role of ethylene in aerenchyma formation, further studies exploring the role of karrikin response pathway in this context are warranted.

Additional insights into regulatory networks underlying aerenchyma formation were provided by the constructed clustered GRN based on the best Arabidopsis hits of identified aerenchyma associated genes. This allowed identification of four hub genes, that are highly connected and thus likely to have a central regulatory role in the network: DNA damage response gene *DRT100,* cell wall hydrolases *XTH16* and *XTH4* and a cell wall protein encoding *AT3G08030.* Interestingly,*DRT100* has been previously implicated in the programmed cell death process of senescence (Ghosh et al., 2013) and was identified as hypoxia responsive in *Prunus* rootstock microarray analysis (Rubio-Cabetas et al., 2018). Likewise, *XTH16* was upregulated by hypoxia in tomato root tissue (Safavi-Rizi et al., 2020); while *X*TH4 is associated with cell wall structure modulation in Arabidopsis (Kushwah et al., 2020) and fluctuates in expression depending on the root location sampled in aerenchyma forming sugarcane roots, with root segments with largest proportion of aerenchyma tissue having highest expression levels of XTH4 (Grandis et al., 2019). Finally, *AT3G08030* is implicated in remodelling of the cell wall and response to a number of stresses in Arabidopsis (Cruz-Valderrama et al., 2019) and has been classified as a molecular marker of seed aging (Garza-Caligaris et al., 2012) but was not previously been connected with waterlogging stress or aerenchyma formation.

Subsequently, analysis of upstream transcriptional regulators of aerenchyma formation was carried out using Arabidopsis homologues of identified aerenchyma associated genes. One of the identified TFs, ERF73 was shown to be induced by hypoxia in *Arabidopsis thaliana*roots and drive ADH expression and metabolic adaptations (Hess et al., 2011). Further, the majority of identified TFs belonged to the ERF (ethylene response factor) family, a class of transcription factors frequently associated with low oxygen responses in numerous species (Hattori et al., 2009; Licausi et al., 2010; Phukan et al., 2018; Xu et al., 2006; Yang, 2014; Yin et al., 2019). As blocking of ethylene perception using 1-MCP partially inhibits aerenchyma formation stimulated by ethylene (Ni et al., 2019), this highlights the importance of ethylene in modulating aerenchyma formation by this phytohormone signalling. TFs with targets overrepresented amongst the aerenchyma candidate genes were previously associated with drought such as DREB2B, DREB19 and MYB94 (Jiang *et al*., 2022; Nakashima *et al*., 2000), salt (e.g. AT1G75490, DREB19 and DREB26) (Akbudak et al., 2018; Kim et al., 2022; Krishnaswamy et al., 2011) and cold stresses (e.g. CRF4, ERF040, ERF57, ERF104 and GATA16) (Illgen et al., 2020; Li et al., 2023; Sun et al., 2016; Zhang et al., 2021; Zwack et al., 2016). While aerenchyma is a key trait for waterlogging tolerance (Mustroph, 2018), it can also be formed in response to other abiotic stresses such as drought (Zhu et al., 2010a), mechanical impedance (He *et al*., 1996) and nutrient deficiency (Postma & Lynch, 2011). Likewise, a recent study has determined the significant overlap of transcriptomic signatures between anaerobic and cold stresses in rice (Thapa et al., 2023), suggesting that similar regulatory pathway may modulate adaptation to these stresses.

The presented work significantly advances our understanding of waterlogging stress induced transcriptional signatures in barley root tissue. Furthermore, exploiting the observed intra-individual variation in aerenchyma formation facilitated isolation of candidate genes associated with this trait and characterization of the regulatory pathways underlying aerenchyma formation. We anticipate that the generated resources will support meta-analyses and future functional studies aimed at development of waterlogging tolerant barley cultivars.

## Author contributions

J.K., P.F.M. and C.K.Y.N. conceived an original idea for the research that O.L.S. further developed. O.L.S., J.O., C.W., F.D. and J.K performed the experiments. O.L.S. carried out statistical analyses with support from R.B., C.W., L.R. and Z.H.. J.K. and O.L.S. drafted the initial version of the manuscript. All authors contributed to the preparation of the manuscript and approved the final version.

## Acknowledgements

We thank Dr. Linda Milne for sending us the barley gene ID conversion file, without which this research would not be possible. The authors thank Prof. Fiona Doohan and Dr. Grace Cott for their comments and suggestions which helped to shape this research. We thank Dr. Graham Hughes for his bioinformatics analysis assistance and comments on the manuscript. We thank the Environmental Protection Agency and Irish Research Council for PhD funding provided to O.L.S. (grant number: irc80800bad9bbf74695ba9c1f3c2982cea). We thank the University College Dublin and the School of Biology and Environmental Science for providing support to the PhD project of R.B. under the PhD Advance Scheme. We recognise SFI Centre for Research Training in Genomics Data Science [18/CRT/6214] for funding of L.R.

## Data availability statement

Data for this article are available as detailed in the article and further requests can be directed to the corresponding author.

## ORCID

Orla L. Sherwood http://orcid.org/0000-0002-8307-902X Rory Burke https://orcid.org/0000-0002-4079-3215 Conor V. Whelan https://orcid.org/0000-0003-1359-930X Frances Downey Louise Ryan https://orcid.org/0000-0002-6876-677X Zixia Huang https://orcid.org/0000-0002-1298-0486 Carl K. Y. Ng https://orcid.org/0000-0001-5832-3265 Paul F. McCabe http://orcid.org/0000-0002-6146-8307 Joanna Kacprzyk http://orcid.org/0000-0003-1054-2367

## Supporting Information

Additional supporting information may be found online in the Supporting Information section at the end of the article.

