## Supplemental Figures, and Supplemental Table Legends for "Transcriptional signatures associated with waterlogging stress responses and aerenchyma formation in barley root tissue"

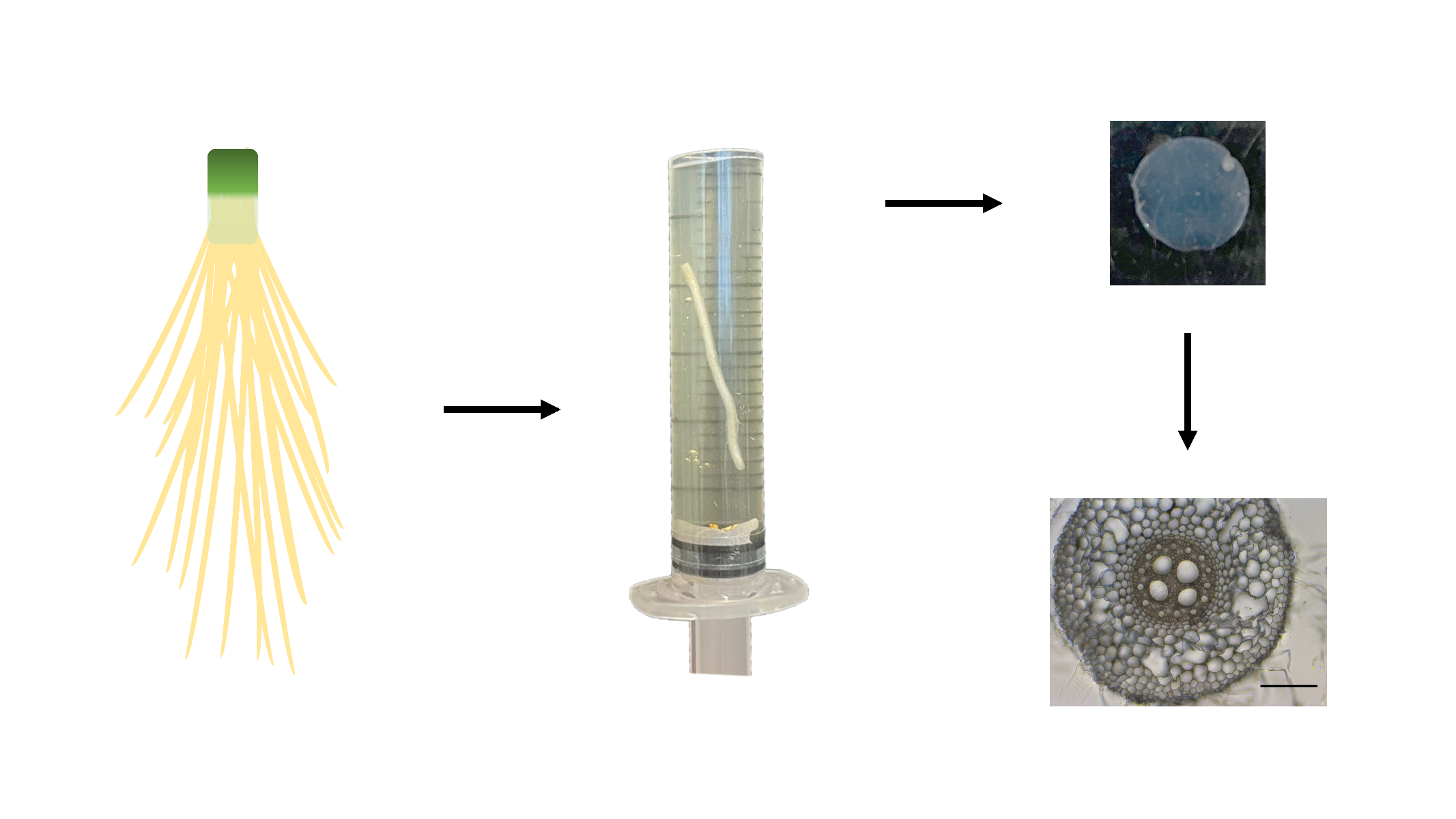

**Figure S1 Hand sectioning experimental set up.** Barley roots were washed and root sections were embedded in liquid agar (5% w/v) inside a 2 ml syringe. Once set, the agar was hand sectioned, mounted on a slide and imaged to examine aerenchyma formation in barley roots. Scale bar = 200 µm.

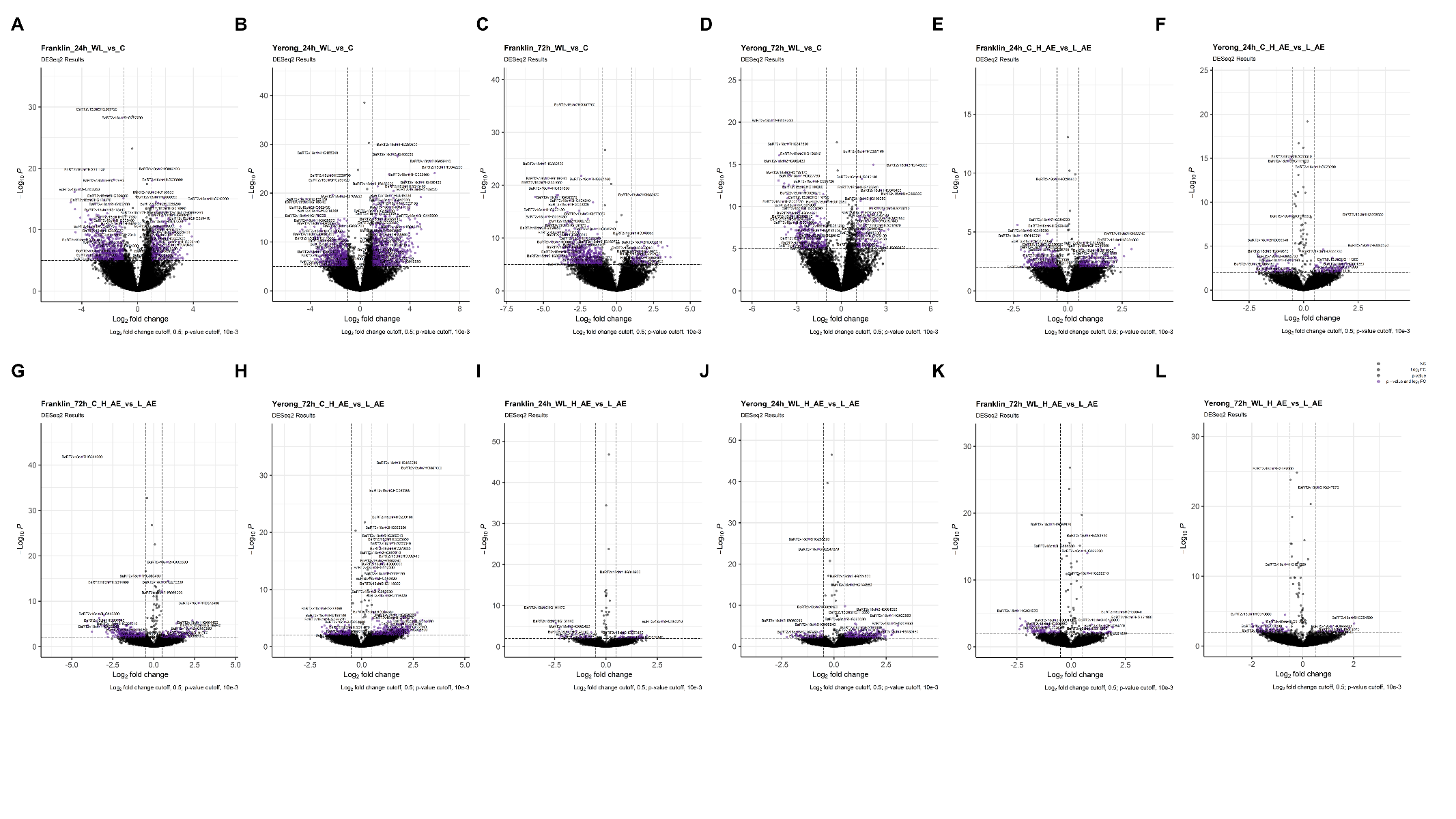
**Figure S2 Volcano plots of DGE analyses.** Volcano plot for each cultivar in WL vs C (**A-D**) and H_AE vs L_AE (**E-L**) comparisons at each timepoint respectively. The second level in the comparisons indicates the reference level. Volcano plots were generated using the R package EnhancedVolcano with log_2_fold change cutoff set to 0.5 and *p*-value cutoff set to 10e-3 (Blighe *et al.,* 2018). C = Control, WL = waterlogged. NS – not significant.

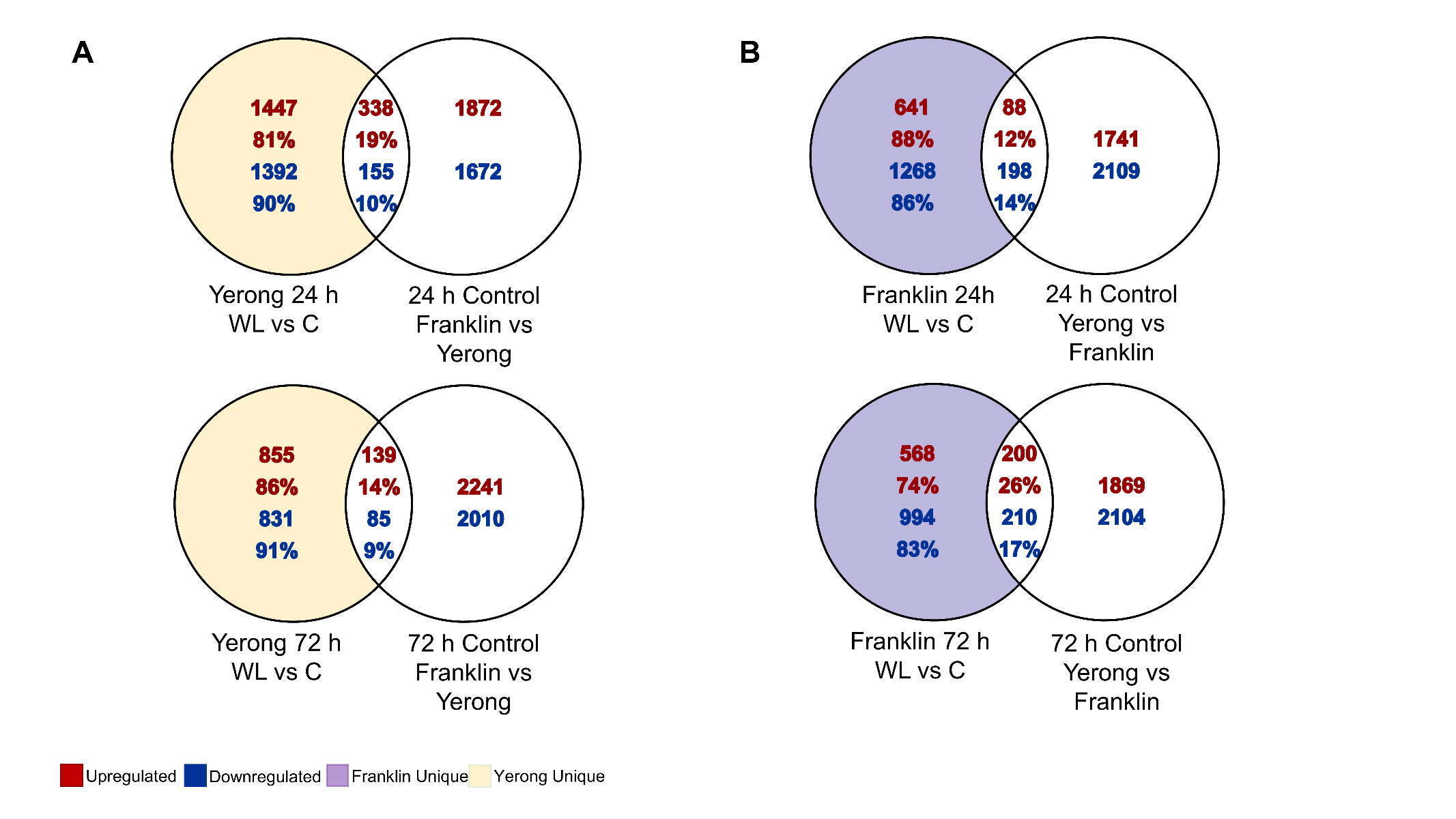

**Figure S3 Basal gene expression levels in barley cultivars compared to waterlogging responsive genes.** Comparison of unique DEGs for each cultivar in WL vs C comparisons are shown compared to the basal gene expression for control treatment between cultivars for 24 h (**A**) and 72 h **(B**) in the Venn diagrams. The second level in the comparison indicates the reference level. Upregulated genes shown in red; downregulated genes are shown in blue. Franklin unique expressed genes are highlighted in purple; Yerong unique genes are shown in yellow areas.

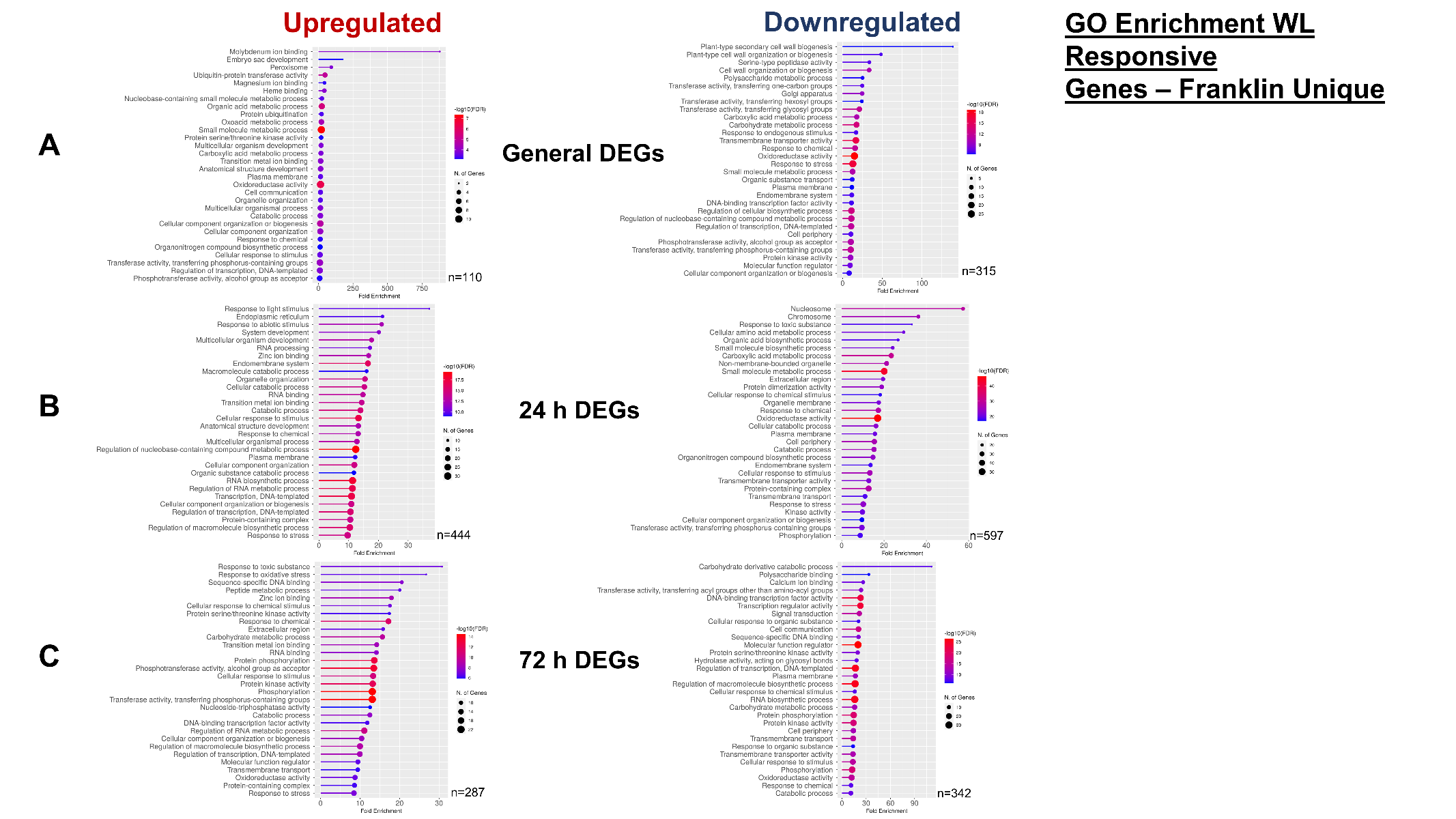

**Figure S4 Gene ontology (GO) enrichment of Franklin WL response genes.** Upregulated and downregulated DEGs unique to the Franklin cultivar for general (both timepoints) (**A**), 24 h unique (**B**) and 72 h unique (**C**) timepoints were identified following DESeq2 analysis. BaRTv2 gene names were converted to MorexV3 gene names (represented by n for each GO analysis) and subsequently input into ShinyGO v0.75c. ShinyGO output was set to 30 GO terms and FDR cutoff as 0.05. The lollipop graph shows fold enrichment on the x-axis with size of dot representing number of genes associated with particular GO term. Colour of line and dot represents -log10(FDR).

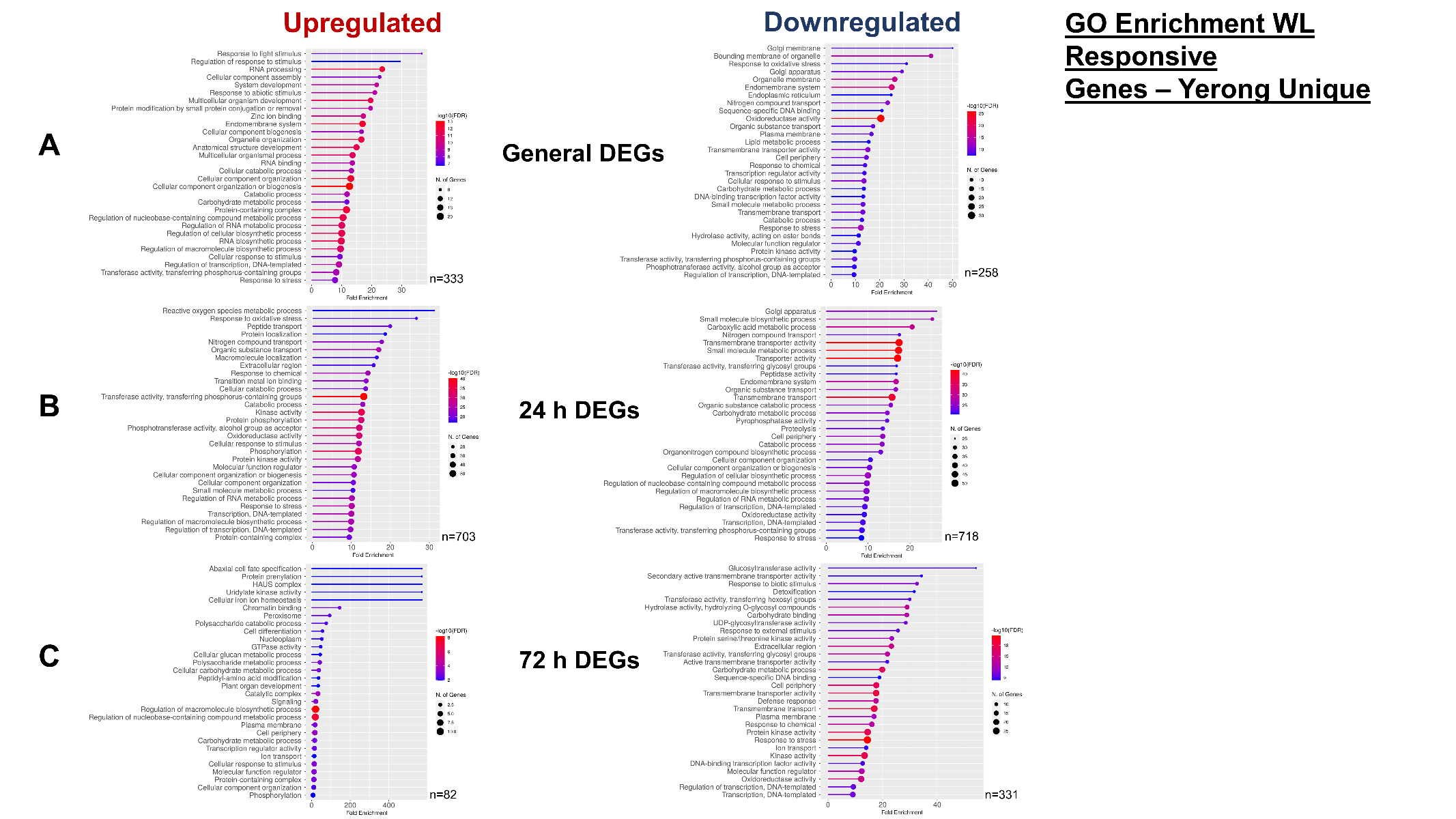

**Figure S5 Gene ontology (GO) enrichment of Yerong WL response genes.** Upregulated and downregulated DEGs unique to the Yerong cultivar for general (both timepoints) (**A**), 24 h unique (**B**) and 72 h unique (**C**) timepoints were identified following DESeq2 analysis. BaRTv2 gene names were converted to MorexV3 gene names (represented by n for each GO analysis) and subsequently input into ShinyGO v0.75c. ShinyGO output was set to 30 GO terms and FDR cutoff as 0.05. The lollipop graph shows fold enrichment on the x-axis with size of dot representing number of genes associated with particular GO term. Colour of line and dot represents -log10(FDR).

**Supplementary Table Legends**

**Table S1. RNA sequencing sample information.** Table presents the sequenced sample information. Pooling information and sample quality parameters are also displayed.

**Table S2. Barley gene ID conversion table.** Table presents MorexV3 IDs with corresponding BaRTv2 IDs.

**Table S3A Transcriptional response of barley roots to 24 h and 72 h of waterlogging stress.** Results of DESeq2 analysis on two barley cultivars and two timepoints of waterlogging stress. Log_2_ fold change of differentially expressed genes is shown for each waterlogging vs control treatment with control set as the reference level for differential gene expression analysis. WL – waterlogged; C – control; NA – no result. **S3B. Differentially expressed genes in barley roots with higher percentage aerenchyma compared with lower percentage aerenchyma.** Results of DESeq2 analysis on two barley cultivars under two timepoints of waterlogging/control treatment. Log_2_ fold change of differentially expressed genes is shown for each high percentage aerenchyma vs low percentage aerenchyma comparison. The low percentage aerenchyma set was used as the reference level for differential gene expression analysis. 81 genes were common to at least two comparison groups of high percentage aerenchyma vs low percentage aerenchyma. WL – waterlogged; C – control; NA – no result. **S3C. Members of aerenchyma associated genes differentially expressed under waterlogging stress treatments.** Aerenchyma-associated DEGs (81) were identified by DGE analysis of high percentage aerenchyma forming roots with low percentage aerenchyma forming roots. These 81 genes were compared with DEGs identified following waterlogging treatment in two cultivars and two timepoints leading to 19 genes common to H_AE vs L_AE comparisons and at least one group of the waterlogging vs control treatment groups. NA – no result.

**Table S4A. Differentially expressed genes in barley roots to 24 h and 72 h of waterlogging stress** compared with Miricescu et al. (2023) hypoxia responsive genes. Using the Miricescu et al. (2023) barley hypoxia responsive gene list after 24 h of waterlogging treatment (98 genes), we obtained a corresponding BaRTv2 gene name where available (total of 46 genes) and compared this dataset with our waterlogging responsive gene lists using the R package GeneOverlap which tests dependency between two lists using a Fisher’s exact test. **S4B. Comparison of waterlogging responsive genes and Miricescu et al. 2023 hypoxia responsive genes.** Using the Miricescu et al. (2023) hypoxia responsive gene list (98 genes), we obtained a corresponding BaRTv2 gene name (46 genes) and compared this dataset with our waterlogging responsive gene lists. Of 46 genes, 44 were differentially expressed in at least one waterlogging vs control comparison. Gene description corresponds to the annotation provided by Miricescu et al. (2023). Likewise, the expression directionality corresponds to the directionality indicated in the Miricescu et al. (2023) dataset. NA – no result.

**Table S5. Aerenchyma-associated gene list transcription factor enrichment.** Aerenchyma associated DEGs were input as BaRTv2 protein FASTA sequences into the ID mapping tool in PlantRegMap to obtain *Arabidopsis thaliana* IDs (n=33). *A. thaliana* gene IDs were used as an input for Transcription Factor Enrichment tool in PlantRegMap (*p*-value cut-off 0.01)***.*** Number(#) gene interactions is the number of genes out of the input gene list which interacted with a specific transcription factor. Q-values are the adjusted p-values found using an optimized FDR approach. Results of a literature search for each transcription factor and its involvement in plant stress responses is represented. NA – no result.
